# Intraspecific trait variability and community assembly in hawkmoths (Lepidoptera: *Sphingidae*) across an elevational gradient in the eastern Himalayas, India

**DOI:** 10.1101/768705

**Authors:** Mansi Mungee, Ramana Athreya

## Abstract

1. Recent progress in functional ecology has advanced our understanding of the role of intraspecific (ITV) and interspecific (STV) trait variation in community assembly across environmental gradients. Studies on plant communities have generally found STV as the main driver of community trait variation, whereas ITV plays an important role in determining species co-existence and community assembly. However, similar studies of faunal taxa, especially invertebrates, are very few in number.
2. We investigated variation of hawkmoth (Lepidoptera: *Sphingidae*) traits along an environmental gradient spanning 2600 m in the eastern Himalayas and its role in community assembly, using the morpho-functional traits of body mass (BM), wing loading (WL) and wing aspect ratio (AR).
3. We employ the recently proposed *T-statistics* to test for non-random assembly of hawkmoth communities and the relative importance of the two opposing forces for trait divergence (internal filters) and convergence (external filters).
4. Community-wide trait-overlap decreased for all three traits with increasing environmental distance, suggesting the presence of elevation specific optimum morphology (i.e. functional response traits). Community weighted mean of BM and AR increased with elevation. Overall, the variation was dominated by species turnover but ITV accounted for 25%, 23% and <1% variability of BM, WL and AR, respectively. T-statistics, which incorporates ITV, revealed that elevational communities had a non-random trait distribution, and that community assembly was dominated by internal filtering throughout the gradient.
5. This study was carried out using easily measurable morpho-traits obtained from calibrated field images of a large number (3301) of individuals. That these also happened to be important environmental response traits resulted in a significant signal in the metrics that we investigated. Such studies of abundant and hyperdiverse invertebrate groups across large environmental gradients should considerably improve our understanding of community assembly processes.

## 1. Introduction

Convergence, and divergence of traits of co-occurring species highlight the effects of two ecological processes that govern community assembly. The abiotic environment, via environmental filtering, causes trait convergence by imposing constraints on the range of trait values, regardless of species, that facilitate their persistence in a habitat (Weiher et al., 1998). On the other hand, competitive interactions are expected to cause trait divergence by limiting the extent of ecological similarity (and hence ‘debilitating’ competition) ‘permissible’ for co-occurring species in a community (Macarthur & Levins, 1967). Consequently, community assembly depends on an interplay between both, biotic interactions and environmental filtering, acting over ecological and evolutionary time-scales.

Several metrics of ‘functional diversity’ (FD) proposed in recent years (Villéger et al., 2008; Mouchet et al., 2010) allow the assessment of the relative importance of these two ecological processes and contributed to an improved understanding of community assembly mechanisms (Albert et al., 2010; Baraloto et al., 2012; Pigot et al., 2016).

However, these metrics, while useful, have ignored several other processes of community assembly such as equalizing fitness or facilitation (Grime, 2006; Butterfield & Callaway; 2013). Additionally, most metrics of FD assume that the relative importance of the two assembly processes is similar across different traits, thereby overlooking the importance of trade-offs in trait filtering (e.g. Spasojevic & Suding; 2012). FD indices also ignore the effects of intraspecific variability (ITV), i.e. they assume that trait variation among individuals of a species is negligible as compared to variation across species. However, this assumption has rarely been empirically validated (but see Albert et al., 2010; Jung et al., 2010) and almost all such attempts have been for plants (but see Griffiths et al., 2016 for a case study on dung beetles) and similar investigations are still lacking for invertebrates which form the bulk of terrestrial biodiversity (see Brosseau et al., 2018; Wong et al., 2019 for recent reviews).

A recently proposed suite of functional trait metrics, *T-statistics* (Violle et al., 2012), incorporate ITV into their calculations. They classified the different processes into the two broad categories of ‘external’ and ‘internal’ filters. While the external filters include all assembly processes outside the community that are responsible for ‘filtering’ species from the regional pool (e.g. environmental constraints, predatory pressures, etc.), the internal filters refer to assembly processes internal to the community, i.e. micro-environmental heterogeneity and density-dependent processes that facilitate coexistence within the community. They used the variance ratios of functional traits across taxonomic (individual, population, species and community) and spatial (local and regional) scales to identify the dominant operational filter.

This identification of two ecological filters, provides a clearer, even if somewhat broad, picture of their relative importance in the establishment and persistence of traits and taxa in the community through the comparison of intra- and interspecific trait variation at local and regional scales (Hulshof et al., 2013; Le Bagousse-Pinguet et al., 2014; Luo et al., 2016; Neyret et al., 2016; Outreman et al., 2017; Xavier Jordani et al., 2019).

The application of this metric to animal taxa has been constrained by the difficulty in identifying the minimum set of traits needed to adequately describe species’ resource axes and in obtaining large scale, individual level trait measurements. Such databases are more easily obtained for plants, which lack behavioral and movement related responses, and where not only the traits, but also their relation to the individuals’ fitness, or their functionality, are easily quantifiable (Lavorel et al., 2013; Lamanna et al., 2014).

We present here the distribution patterns of three key morphological traits – body mass (BM), wing loading (WL) and wing aspect ratio (AR) – of 3301 individual hawkmoths (Lepidoptera: *Sphingidae*) across a large environmental gradient at multiple taxonomic and spatial scales. We have analysed these patterns to assess (i) the functional role of these traits and (ii) the relative importance of external (e.g. environment) and internal (e.g. competition) filters, vis-a-vis randomness, in structuring elevational communities. The environmental gradient was derived by a linear combination of temperature, precipitation, air density and productivity across 2600 m of elevation in the eastern Himalayas.

More specifically, we test the following hypotheses –

i. The degree of overlap of the distribution of traits across communities should be anti-correlated with the environmental distance between them as the three traits have been previously linked to organismal performance (Sinervo & Huey, 1990; Hammond et al., 2000; Berwaerts et al., 2002; Huyghe et al., 2005; McGill et al., 2006) through thermoregulation and flight (Dudley, 2002; Dillon et al., 2006)
ii. The community mean trait values of (a) BM should increase with elevation (better thermoregulation associated with larger bodies), (b) WL should decrease and AR should increase with elevation (improved flight efficiency at higher elevations where resources are generally sparse and air density is lower),
iii. If hawkmoth communities are governed by the same widespread pattern seen in other taxa (Seifert et al., 2015), the change in community mean trait along the environmental gradient will be dominated by species turnover rather than intraspecific variation, and
iv. Hawkmoth communities at each elevation must be non-random subsets of the regional pool and, internal filters should dominate community trait structure.

## 2. Methods & Materials

### Study area and Field Sampling

Hawkmoth sampling was carried out in Eaglenest Wildlife Sanctuary (hereafter EWS), a Protected Area of 218 km^2^ located between 27° 02” 09’ N and 92°18” 35’ E in the eastern Himalayas of Arunachal Pradesh, northeast India, during April – July 2014 (**Figure 1**). The high diversity of this region, believed to be due to its complex terrain and its location at the confluence of the Oriental and Sino-Japanese floristic and faunistic zones (Holt et al., 2013) makes it a globally important biodiversity hotspot (Orme et al., 2005). The large altitudinal range of 3000 m coupled with high rainfall (> 3000 mm along the southern slopes) has resulted in diverse habitat types ranging from tropical wet evergreen below 900 m to coniferous temperate forests above 2800 m (Choudhury, 2003). The sampling, in the form of point surveys at UV illuminated screens on no-moon nights, was carried out along a vehicle track characterized by roadside scrub in very close proximity to primary forest (5-20 m away). The 12 elevations between 500 m and 2800 m were clustered in a small stretch spanning just 15 km. The 200 m location, near the village of Tippi, was separated from its nearest neighbor by about 20 km due to the lack of access to suitable habitat along this road (**Figure 1**).

**Figure 1.**
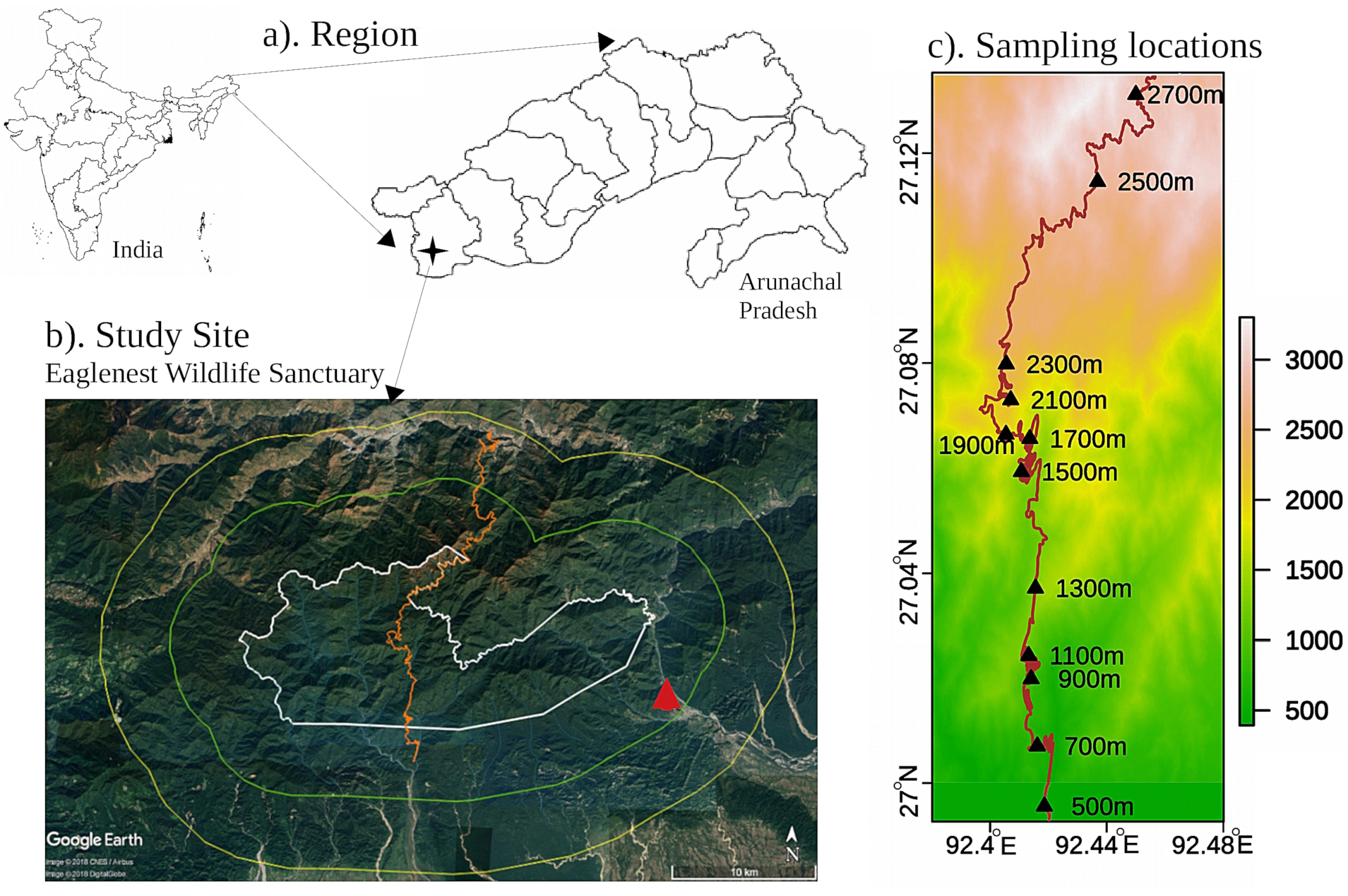
Study site in Eaglenest wildlife sanctuary, India. a). Location of the study site in West Kameng district, Arunachal Pradesh, north-east India b). A google earth image of Eaglenest Wildlife Sanctuary with the the boundary of the Protected Area (PA) marked in white, and the 5 km buffer strip in green. The dirt track running through the sanctuary traverses elevations from 100 m in the south to the pass at 2780 m and down to 1200 m to the north. is shown in orange, and the 200 m sampling location, which was outside the PA is marked by a red triangle. c). Digital elevation map showing the Eaglenest track and the sampling locations between 500 m and 2700 m. The 200 m sampling site is shown by a red triangle in the Google image.

The sampling was limited to a single compact transect to minimize the bias due to a variation in gamma diversity while sampling across distant transects (McCain, 2007). We sampled at 2-5 elevations simultaneously to sample across the elevational gradient with some degree of uniformity of weather conditions (which can change drastically from day to day). Hawkmoth individuals arriving at a light screen were photographed against the reference grid (on the screen) using consumer-grade point-and-shoot digital cameras. Following Willott (2001) our sampling strategy attempted to equalize the number of individuals, rather than trap nights, across elevations to minimize the large diurnal variation in moth numbers at a light screen, even within the no-moon window. A total of 4731 hawkmoth individuals, spanning all 3 Sphingid subfamilies, 30 genera and 80 morphospecies, were recorded from across 13 elevations.

### Species identification and trait measurement

We assigned individuals to morpho-species using the online resources made available by Kitching and collaborators (e.g. Kitching, 2019). We obtained the primary measurements of body length, thorax width, wing costum length and wing breadth from which we derived the three functional traits of body mass (BM), wing loading (WL) and wing aspect ratio (AR) from field images after calibration and distortion corrections (Mungee & Athreya, 2019). Traits were reliably measured for 3301 images (69% individuals) spanning 76 morphospecies and 30 genera making it the first systematic compilation of any insect trait data from the region. Details of the sampling methodology and trait measurement are provided as Supporting information <u>**(Supplementary S1)**</u>.

### Environmental variables

We used 4 environmental variables including mean annual temperature (MAT), mean annual precipitation (APPT), productivity (EVI: enhanced vegetation index) and air density (AD). These variables were strongly correlated with each other and all decreased with elevation. A principal component analysis showed that the first two axes explained 91.4 % and 7.7 % of the variance. Therefore, we used PC-1, which has a strong linear relationship with elevation, as a composite environmental variable, hereafter referred to as the *Environment* in italics (see <u>**Supplementary S1**</u> for a detailed analysis).

### Diversity and Trait data set

Traits were measured for only a subset (trait data set: 3301 individuals; 69%) of the total number of individuals identified to morphospecies (diversity data set: 4731 individuals). We carried out all analyses using just the trait data set, and also with the diversity data set; in the latter we filled in the missing traits by randomly drawing trait values from other individuals of that species at that elevation. For example, only 66 of the 79 individuals of *Acosmerycoides harterti* at 700 m had measured traits. The remaining 13 individuals were assigned trait values drawn from the set of 66 measurements (random sampling with replacement). We did not simulate these extra values using the population mean and standard deviations as this would have affected the statistics of the sample mean and dispersion. The results for both treatments are quite similar. In any case, an assessment of the completeness of our samples using taxonomic rarefaction (for diversity data set) and functional rarefaction (for the trait data set) curves was done (<u>**Supplementary S1; Figures 6 & 7 respectively**</u>). Additionally, since the fraction of individuals with traits was very low at 1700 m (due to poor weather), we carried out the analysis with and without this elevation.

The analysis with the trait data set including 1700 m is shown here; the same analysis with (i) trait data set without 1700 m and (ii) diversity data set with 1700 m are shown in <u>**Supplementary S2**</u>.

#### Trait variation across the environmental gradient

We used two approaches to examine the functional response of hawkmoth communities across the environmental gradient. First, we investigated the change of functional ‘alpha’ diversity across the gradient using the community abundance-weighted mean trait value (CWM; Lavorel et al., 2008). Two different CWM values were analysed for each community, CWM1 – using the regional species mean trait value, i.e. mean across all elevations, and CWM2 – using the population mean trait values, i.e. mean across all individuals of a species at an elevation:

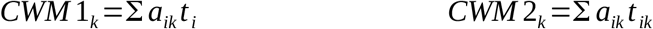

where *a*_*ik*_ is the relative abundance of species *i* at elevation *k*, *t*_*i*_ is the regional mean trait value for species i, and *t*_*ik*_ is the population mean for species *i* at elevation *k*. The change of CWMs with environment was assessed using the ordinary least squares (OLS) regression.

Second, we quantified the change in trait across the gradient by the degree of overlap of the kernel density distributions for all pairs of communities (Mouillot et al., 2005), i.e essentially the functional ‘beta’ diversity. The kernel density distributions were constructed in a non-parametric manner and do not assume an underlying distribution for community trait values. The distribution function for each trait at each elevation was calculated as the sum of kernel density functions for each individual in that community. The degree of overlap of a trait between any pair of communities was simply the area of overlap of their trait density distributions. We used OLS regressions to examine change of overlap with environmental distance. We did not estimate the multivariate overlap as either the average overlap across all traits, or the overlap in multidimensional space (Mouillot et al., 2005). The difference in the slopes for different traits, i.e. the rate of response to the same environmental gradient, can be a measure of the strength of their functional response.

#### Variance decomposition

We partitioned the community-level response of hawkmoth traits to the environmental gradient into species turnover (STV) and intraspecific variation (ITV) following the approach of *Lepš* et al., (2011).

#### Trait robustness

We used linear discriminant analysis (LDA; Venables & Ripley, 2002) to assess if the hawkmoth species were well discriminated by the four primary traits (body length, thorax width, wing costum length and wing breadth). This helped to quantify the role of intraspecific trait variation in confounding the assignment of an individual to its species on the basis of (just) these four traits.

#### T-statistics

We first log-transformed the trait values to remove potential scaling effects between measurements. We calculated three variance ratios (*T-statistics*, Violle et al., 2012) at nested spatial and taxonomic scales as follows:

- 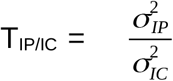, where σ^2^_IP_ is the variance of trait values among individuals within a population, and σ^2^_IC_ is the variance of trait values among individuals within a community (strength of internal filtering).
- 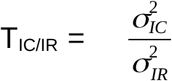, where σ^2^_IR_ is the variance of trait values among individuals within the regional pool (strength of external filtering acting on individuals)
- 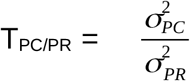, where, σ^2^_PC_ is the variance of population mean trait values within a community and σ^2^_PR_ is the variance of population mean trait values within the regional pool (strength of internal filtering acting on species)

The observed metrics were compared to those obtained from the simulated null models to detect non-random assembly of community traits. Details on generation of the null models are provided in <u>**Supplementary S3**</u>. The standardized effect sizes (SES) were calculated as:

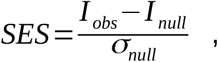

where *I*_*obs*_ is the observed value for a given index, *I*_*null*_ is the mean of the simulated null replicates, and *σ*_*null*_ their standard deviation. OLS regressions were used to assess the relationship between individual metrics and the composite environmental variable and also for T_IP/IC_ *versus* species richness (Violle et al., 2012).

All above analyses were performed in the R programming software; version 3.4.4 (R Development Core Team, 2015) and the following packages were used: *vegan 2.5.4* for computing species richness, diversity indices, taxonomic rarefaction curves and environmental variables PCA scores (Oksanen et al., 2019); *evolqg 0.2.6* for functional rarefaction curves (Melo et al., 2015); *FD 1.0.12* for CWM analysis (Laliberté et al., 2010); *sfsmisc 1.1-3* for the trait kernel density analysis (Maechler, 2019); *MASS 7.3.51.4* for LDA (Venables & Ripley, 2002) and *corrplot 0.84* for the associated plot (Wei & Simko, 2017); *cati 0.99.2* for calculating the *T-statistics and* generating null models; function *decompCTRE* for variance partitioning (Taudiere & Violle, 2016).

## 3. Results

### Trait variation across the environmental gradient

Community weighted mean of BM exhibited a significant positive relationship with *Environment* (**Figure 2**). The slope of CWM2, that incorporates intraspecific variation in it’s calculation was slightly greater than CWM1. WL did not exhibit any significant relationship with *Environment* using either CWM1 or CWM2. CWM of AR showed a significant positive correlation with *Environment* and similar to BM, the slope of CWM2, was marginally greater than CWM1 **(Table 1)**. We used Fisher’s r-to-z transformation to compare the two fits. The difference between the slopes of CWM1 and CWM2 were not significant for any trait <u>**(Supplementary S3; Table 2)**</u>. LDA analysis yielded a mean attrition probability of 63% for correctly attributing an individual to its species from its trait values (<u>**Supplementary S3**</u>, **Figure 1**).

**Table 1.**
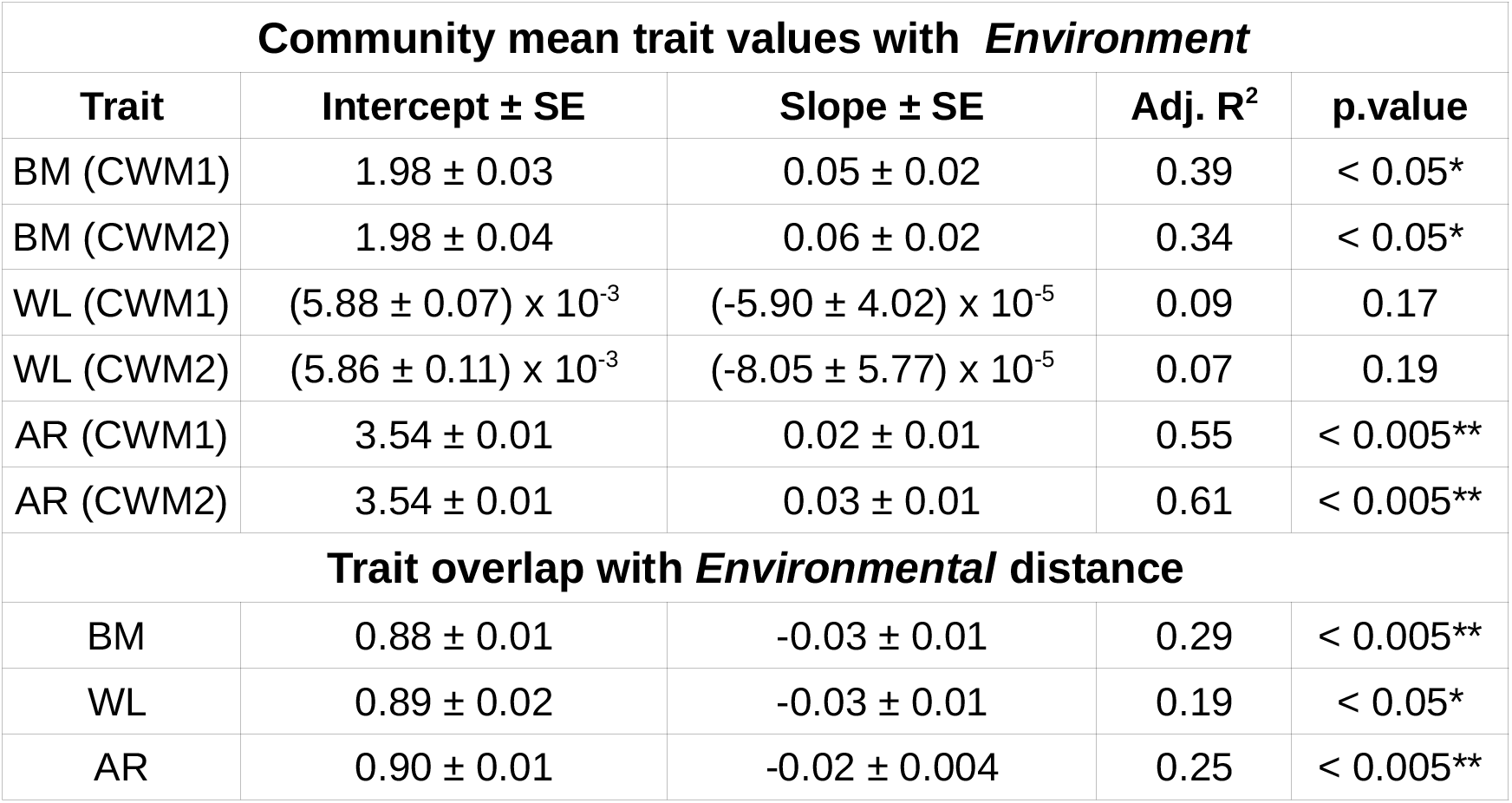
Variation of hawkmoth community traits with *Environment*. Results from the linear regression analysis involving community traits and the Environment (i.e. composite environmental variable, which is also strongly correlated with elevation). The traits used are body mass (BM), wing loading (WL) and wing aspect ratio (AR). The analysis was carried out for both population mean trait values weighted by local abundance (CWM1) and regional species mean trait values weighted by local abundance (CWM2).

**Figure 2.**
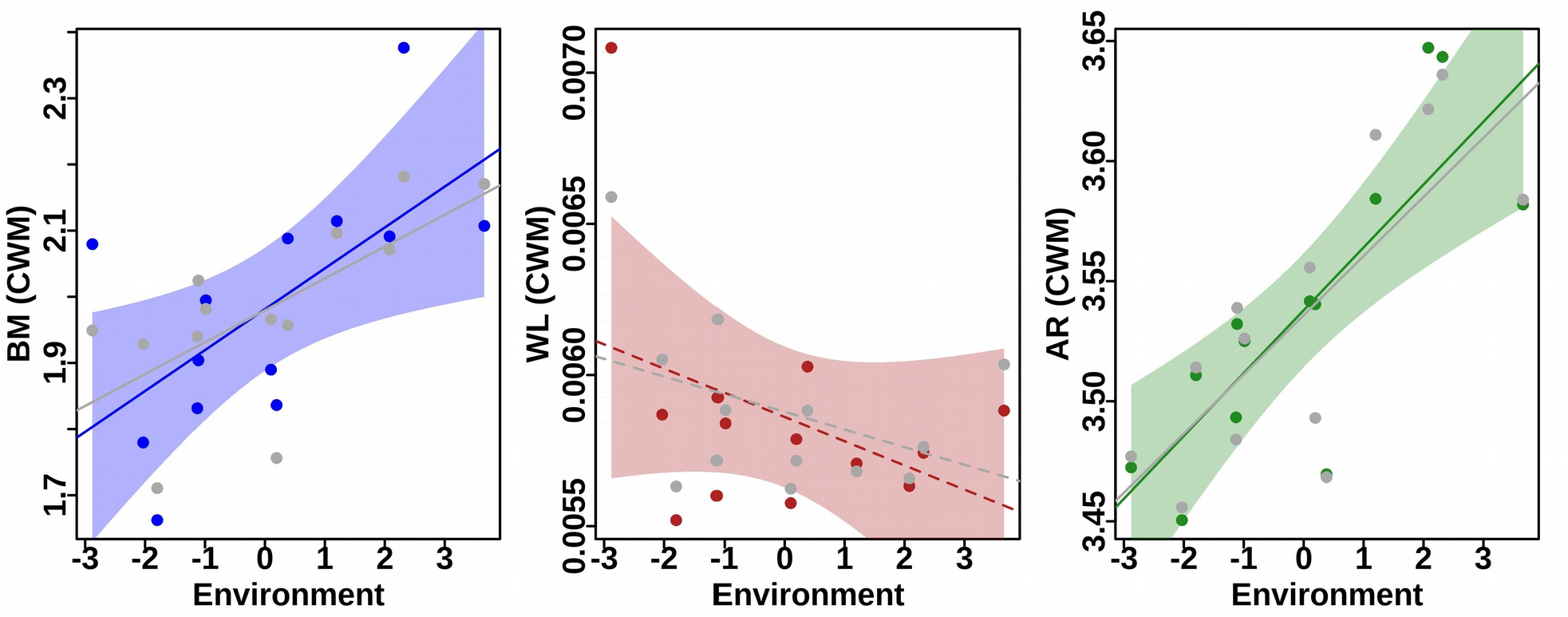
Variation of hawkmoth community traits along an *Environment*. The plots show the change in community weighted means (CWM) of body mass (BM), wing loading (WL) and wing aspect ratio (AR) plotted against composite E*nvironment* (which has a strong positive correlation with elevation). For each trait, the colored representations are for CWM1 (population mean trait values weighted by local abundance) while the grey are for CWM2 (regional species mean trait values weighted by local abundance). The lines represent the best fits, with the ones significant at 95% shown in solid and the others with dashes.

Kernel density plots for each trait-elevation distribution are shown in <u>**Supplementary S3;**</u> **Figure 2**. The reduction of trait overlap with environmental distance **(Figure 3)** was significant for all traits **(Table 1)**. Slopes were significantly different between BM and AR (Fisher’s r-to-z transformations, <u>**Supplementary S3; Table 3**</u>). Trait overlap did not exhibit any significant pattern with absolute elevation, for any of the three traits <u>**(Supplementary S3; Table 4)**</u>.

**Figure 3.**
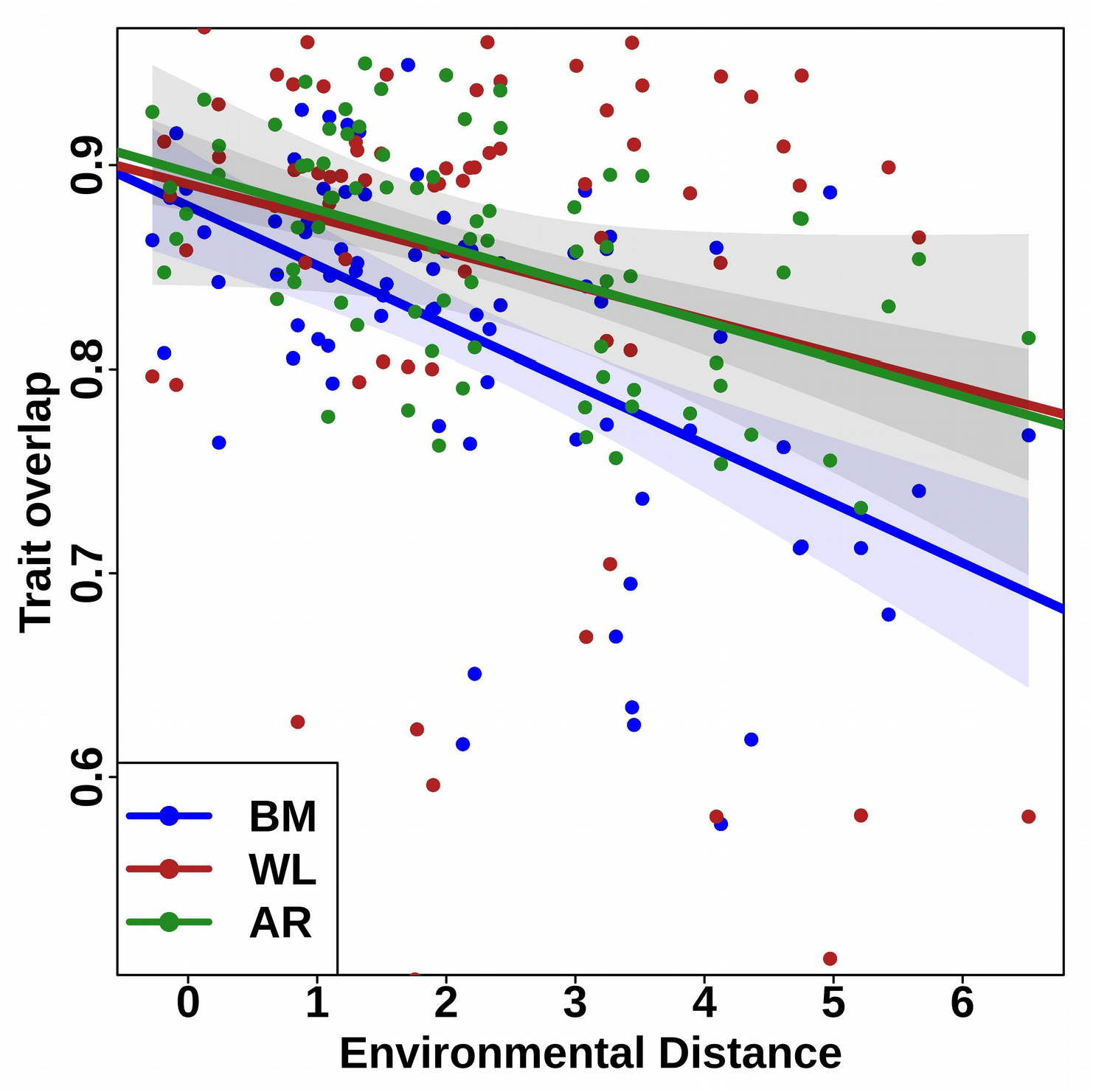
Relationship between trait overlap and environmental distance. The plot shows the scatter and the regression line for the relationship between the overlap in trait distribution functions for pairs of communities and the *Environment*al distance between them. The shaded areas are the 95% errors on the slope. The three traits analysed are body mass (BM), wing loading (WL), and wing aspect ratio (AR). The statistically significant correlation suggests the importance of these as key functional response traits.

### Variance decomposition

The maximum source of variation in all three traits was species turnover (BM = 70%, WL = 50%, and AR = 99%), followed by intraspecific variation (BM = 25%, WL = 23% & AR = <1%; **Figure 4**). This was also reflected in the high species turnover across the entire gradient (<u>**Supplementary S3;**</u> **Figure 3**)

**Figure 4.**
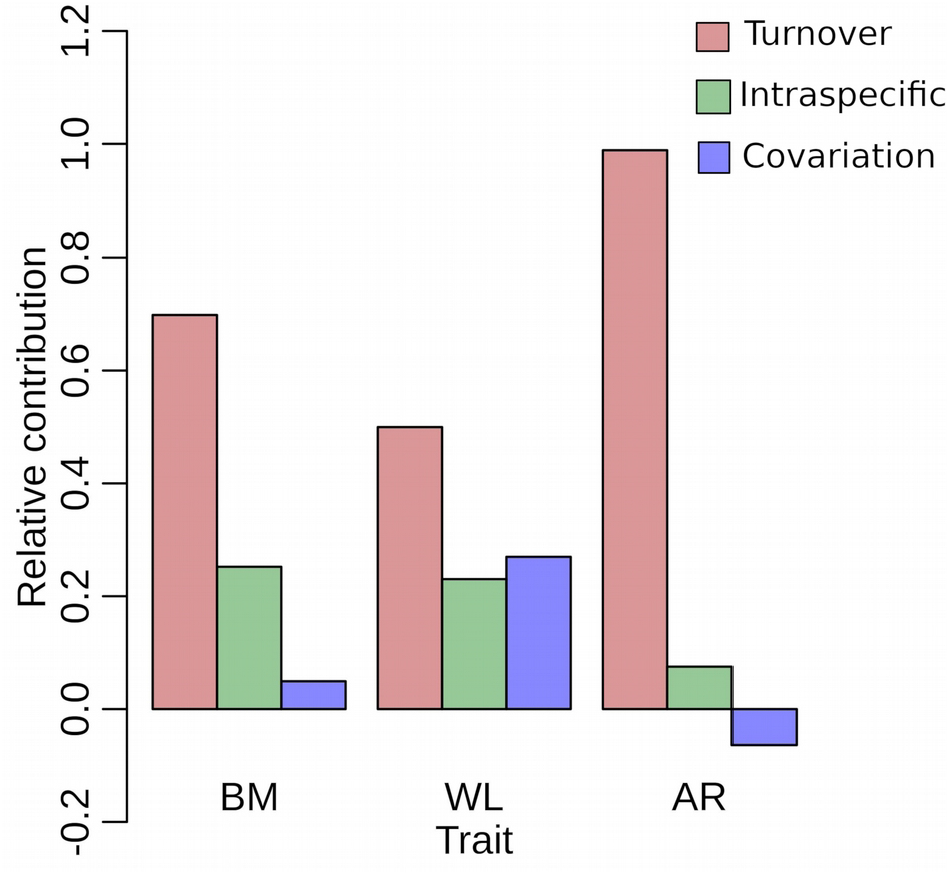
Decomposition of variation of hawkmoth community functional traits along an Environmental gradient. The contributions of species turnover and intraspecific variation (and their covariance) were calculated for body mass (BM), wing loading (WL) and wing aspect ratio (AR) using the approach of *Lepš* et al., (2011).

### T-statistics

The observed metrics of *T-statistics* and their SES values are listed in <u>**Supplementary S3; Tables 5-8**</u>. T_IP/IC_ was significantly lower than the null model at all elevations and for all three traits (**Figure 5**). Distribution of T_IC/IR_ was more variable: BM was significantly lower than null at some of the lowest (200, 500, 900 m) and highest (2500 & 2700 m) elevations and higher than null in between (1100, 1300, 1500, and 1700 m). SES for T_IC/IR_ value for WL was significantly lower than null at most elevations, but higher at 200 m. AR was mostly not significantly different from null, except at 200 m (lower) and 700 m (higher). T_PC/PR_ was not significantly different from null at any elevation, and for any trait (**Figure 5**). We also observed that the correlation of SES values with *Environment* was significantly negative for T_IP/IC_ of BM and WL, and significantly positive for T_IC/IR_ of WL. T_IP/IC_ was not found to be significantly correlated with rarefied species richness for any of the three traits <u>**(Supplementary S3;**</u> **Figure 4)**.

**Figure 5.**
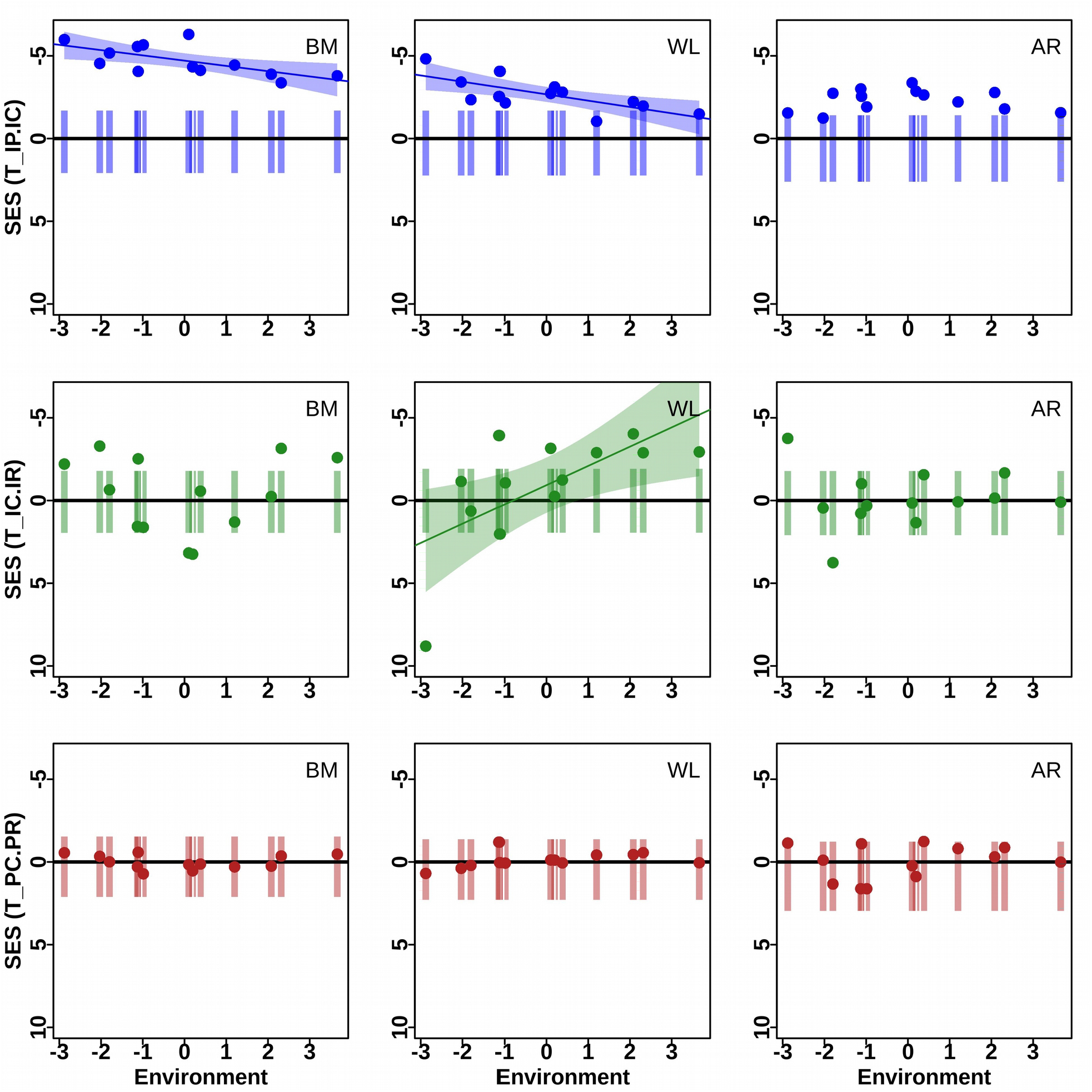
T-statistics of hawkmoth functional traits across an *Environmental* gradient. The plots show the T-statistics for 3 traits: body mass (BM), wing loading (WL) and wing aspect ratio (AR), and for each of the 13 elevational communities (along the x-axis) in terms of the standardized effect size (SES) along the y-axis. The vertical bars are the 95% distribution from 999 simulated null communities, and the dots are the observed community values. The three T-statistics variance ratios are (a) T_IP/IC_ — within-population to within-community; (b) T_IC/IR_— within-community to within-regional, assessed using individual values; and (c) T_PC/PR_— within-community-wide to within-region, assessed using population-means. Additionally, linear regression fits including errors (shaded region) have been shown for metrics which are significantly correlated (CI > 95%) with the composite *Environment*al variable (which is positively correlated with elevation).

## 4. Discussion

This study involves the first systematic collection of individual-level trait data for any invertebrate group from the study region. We are not aware of any previous work that uses trait variance ratios via *T-statistics* to explore community assembly of invertebrate fauna over a continuous gradient and across multiple sites (but see Outreman et al., 2017). This work also highlights the importance of morphological traits as key functional ‘responses’ when the traits can be directly implicated in individual survival, or performance strategies.

We tested the hypothesis that tropical hawkmoth communities of eastern Himalayas are not randomly assembled with respect to key morphological traits. We demonstrated that body and wing sizes are important functional attributes that respond to a changing environment across a large elevational gradient, and the strength of this response did not change significantly upon incorporation of intraspecific variability. Community mean body mass and aspect ratio increased with *Environment* (i.e. elevation), however, wing loading did not exhibit any directional variation along the gradient. Using a trait data from 3301 individuals, we showed strong internal filtering in hawkmoth communities, which indicates low niche overlap among co-occurring species. The strength of the external filtering varied with trait and environment. More importantly, external filtering acted on individuals rather than species highlighting the importance of incorporating intraspecific variance in understanding community assembly processes.

### Trait variation across the environmental gradient

Several frameworks and guidelines have been suggested for consideration in trait selection for faunal groups (for most recent reviews see Luck et al. 2012 for vertebrate groups and Brosseau et al., 2018 & Wong et al., 2019 for invertebrates).

Morphological traits are easily quantifiable, and some can be directly linked to individual function and survival, especially when the performance trait has a simple mechanical basis in design (Wainwright, 1988). Especially for hawkmoths, which are a facultatively endothermic group (thermoregulation by active shivering of thoracic muscles; Heinrich 1996), body mass can be considered a key response trait determining species’ functional strategies along a wide elevation (temperature) gradient. It should be noted that while body mass may be governed by multiple life history traits (e.g. reproductive age, starvation, etc), body volume (our measurements) will have a more predictable relationship with thorax muscle mass. Wing loading and wing aspect ratio are two important determinants of flight performance, and consequently fitness (e.g. reproduction via mate-locating, host-plant searching, and survival via predator escape, resource acquisition and dispersal).

A larger body size at low ambient temperatures is consistent with better thermoregulatory properties for facultatively endothermic hawkmoths, *sensu lato* Bergmann’s Pattern (Salewski & Watt, 2017). Previously reported patterns of variation in body size for invertebrates along temperature gradients have been idiosyncratic to taxon and spatial scale of investigation (Shelomi, 2012; Vinarski, 2014; Brehm et al., 2019). Body size also has direct implications for several important processes such as physiological, macro-ecological and evolutionary (Blackburn & Gaston, 1994). Thus further tests of the relative importance of these different causative mechanisms will be required to ascribe the strong observed pattern to a definite process.

The adaptations for powered flight can be broken down into two key wing attributes – (i) ratio of body mass to wing area (wing loading), and (ii) functional variations in wing shape, especially in the length of the wing relative to the width (aspect ratio) (Hassall, 2015). Our results indicate that hawkmoth communities exhibit strong adaptations for better dispersal abilities and flight efficiency via increased aspect ratio, but not wing-loading. In vertebrates, higher aspect ratio (longer, thinner wings) is found to give faster and more efficient flight and has been shown to be associated with migratory species in birds (Vágási et al., 2016). For insects, while there are several theoretical speculations that lower aspect ratio may be more suited at higher air densities (due to higher viscous forces experienced by small objects, and due to the difference in the number, structure and locomotory independence of wings between insects and higher vertebrates), empirical results have been mixed (Hassall, 2015; Bai et al., 2016). A higher aspect ratio for communities of hawkmoths at higher elevations, is indicative of adaptations to long dispersal flights which may be better suited to the patchiness in resource distributions more commonly associated with regions of low productivity, as suggested by the EVI values of high elevations in our data. Interestingly, wing loading did not exhibit a significant trend with elevation indicating different selection pressures on wing shape and size.

Trait overlap decreased with increasing *Environmental* distance between communities confirming the role of the three morphological traits in functional response strategies of the hawkmoth communities along an elevational gradient that exhibits a large range of temperature, productivity and air density. Many studies have demonstrated correspondence between species morphological traits (morphospace) and their ‘performance’ or functional strategies (Price et al., 2014; Pigot et al., 2015; Dehling et al., 2016). For instance, Pigot et al. (2015) found that key dimensions of the ecological niche in passerines, including diet, foraging maneuver and foraging substrate were, to varying extents, predictable on the basis of morphological traits. As agents of many ecosystem processes and services, the morphological response of non-producers merits further research.

### Variance Partitioning

Species turnover and intraspecific variation accounted for 73% and 19%, respectively, of the total trait variance. This not insignificant contribution of ITV is consistent with a growing body of literature advocating the use of both individual and species-specific traits to investigate community assembly mechanisms (Jung et al., 2010). As seen in plant communities, the contribution of intraspecific variation was strongly dependent on the trait, with 25% contribution for BM, and less than 1% for AR. Very few studies have explored the extent of intraspecific variability in insect communities till date, and obtained very contrasting values (< 5% for dung beetles, *Insecta: Coleoptera,* by Griffiths et al., 2016; < 1% for stonefly assemblages, *Insecta: Plecoptera*, by Garcia-Raventós et al., 2017; and > 70% for spiders, Arachnida: Araneae, by Dahirel et al., 2017). Other taxa have reported mixed results (33% for tadpoles by Xavier Jordani et al., 2019; 70% in lichens by Asplund & Wardle, 2014). In general, it is suggested that community-level trait variation, especially across broad environmental gradients, is driven primarily by species turnover, but the relative importance of intraspecific variation depends strongly on the trait, environmental factor, and spatial scale considered. The variation observed across studies for invertebrates warrants further investigations to propose general assembly mechanisms.

We note here that while intraspecific variability played a lesser role in the change of the community mean trait value with *Environment*, the LDA analysis of species-trait robustness <u>**(Supplementary S3**</u>, **Figure 1)** suggests a substantial spread of values away from the species mean.

### Internal and external filtering

The realized and fundamental niches of co-occurring species, are key to understanding how local communities are assembled from a ‘regional’ species pool (Kraft et al., 2008). Using *T-statistics*, we demonstrated the non-random assembly of hawkmoth communities and classify them based on their assembly forces i.e. internal and external filtering. The metric T_IP/IC_, indicative of internal filtering, showed that individuals of a species within a local community were more closely clustered with each other in trait space, than may be expected from a random chance. Similar patterns of non-overlapping distribution of morphological traits among co-occurring species has been reported previously for tadpoles (Xavier Jordani et al., 2019) and for plant communities across environmental gradients (Luo et al., 2016; Neyret et al., 2016). Interestingly, the SES values of TIP/IC exhibited a negative relationship with *Environment* for body mass and wing loading, suggesting a decrease in the strength of internal filtering with elevation (i.e. increasing environmental severity).

The results were more variable for the metric T_IC/IR_. Values for BM were lower than null at some of the highest and lowest elevations, but those for WL were consistently lower than null for most communities (i.e. individuals within a community are more similar to each other than individuals across communities), indicative of external filtering. External pressures, perhaps the stressful environment (low productivity, temperature and air density at high elevations) and anthropogenic or predator pressures (at low elevations), are acting as a strong filter allowing only individuals with specific trait values to persist in these communities. Values higher than the null distribution for certain communities is a potentially interesting result and indicates that the individuals here exhibit high variance in their trait values, compared to the variance in trait values of individuals across all communities and thus, these communities are assembled with weak external filtering. Importantly, similar patterns of external filtering were not observed at the species level (T_PC/PR_), when only interspecific variation was used. This supports the importance of incorporating intraspecific variation in examination of community assembly patterns (Jung et al., 2010).

## CONCLUSION

We carried out a study of intraspecific traits of the hawkmoth community across a 2600 m elevational transect in Eaglenest wildlife sanctuary in north-east India. We obtained a diversity data set of 4731 individuals spanning 80 species and 30 genera, of which we could measure traits for 3301 individuals. This is the first such systematic study of intraspecific variation of any taxon from this globally important biodiversity hotspot. We found that the three traits – body mass, wing loading and wing aspect ratio, which are implicated in thermoregulation and flight – change in response to a changing environment across that elevational gradient. As a community, hawkmoths exhibited larger body sizes, lower wing loading and higher aspect ratio at higher elevations. Species turn-over dominated these changes but the intraspecific variation was not insignificant; furthermore, its contribution changed with the trait. We also used a metric from the suite of T-statistics to infer the role of internal filtering in community assembly. Although changes in community-average trait values across the environmental gradient may be discerned by (regional) species-level means, as has been the dominant strategy of investigations, one can/should expect local community dynamics to be largely influenced by intraspecific trait variation. Indeed, another T-statistic metric showed a significant signal only when individual trait values were used but not with species means. Only plant researchers seem to have used T-statistics to investigate internal and external filters affecting community assembly, except for two, recent, studies on hymenopterans (Outreman et al., 2017) and tadpoles (Xavier Jordani et al., 2019). Thus, this study is perhaps the first such of a group from the hyperdiverse Lepidoptera.

## Acknowledgments

The authors would like to thank Deepak Barua for stimulating our interest in functional ecology and for the several rounds of discussions that led to the synthesis of this paper. We thank the Forest Department of Arunachal Pradesh for their assistance and research permits (CWL/G/13(95)/2011-12/Pt.II/660-62 during 2011-2015). This work would not have been possible without the enormous support of diverse kinds provided by Mr. Indi Glow, Nima Tsering and other members of the Singchung Bugun community.

