## Supplementary_01 for "Intraspecific trait variability and community assembly in hawkmoths (Lepidoptera: *Sphingidae*) across an elevational gradient in the eastern Himalayas, India"

### Hawkmoth Sampling

Hawkmoths were sampled at UV light screens, during April – July, 2014, set up along a vehicle (dirt) track traversing Eaglenest wildlife sanctuary between 400 m and 2800 m. Portable light screens were set up at 2-5 elevations between 7 PM and midnight during the "no-moon" phase; i.e. during the 7 days before and 3 days after the new moon when the moon was below the visible horizon between 7 PM and midnight.

Hawkmoths which arrived at the screen were photographed, then captured for marking (by clipping a tiny portion of the forewing apex) to avoid double counting, and for collection of the two middle legs for DNA, and subsequently released.

The field strategy was to collect similar number of individuals at each elevation because of the high daily variability observed in hawkmoth numbers at a light screen, even within the 10-day no-moon period (See Figure 1 below). Previous studies have also reported high fluctuations in moth activity due to local weather, temperature, wind, cloud, rains, etc (McGeachie 1989, Schulze & Fiedler 2003, Beck et al. 2008), and argued that the number of individuals is a better measure of the sampling effort for moths than the number of nights (Willott 2001). In particular, we observed flushes of individuals of some species on many nights which were clearly non-Poissonian events.

We recorded a total of 4808 hawkmoth individuals, of which 4,731 could be reliably identified to a morpho-species. We refer to this as the "diversity data set" which was used in all analyses of taxonomic diversity (e.g. beta diversity, species richness, etc.). We were able to measure traits for a subset of the diversity data, the "trait data set" of 3,301 individuals. We could not measure traits of those individuals which did not sit on the reference grid, and for a small number of images in which the distortion could not be corrected.

The results are summarized in **Tables 1 and 2, and Figure 1**

### Hawkmoth Trait Data

We obtained shape and size calibrated images by photographing moths against a reference rectangular grid on the fabric of the UV light screen. The distortion of this background rectangular grid (of known dimensions) were used to calibrate the image for shape and size.

We measured the following traits on a calibrated image using the 8 landmarks A-H shown in Figure 2:

1. Body: Body length = $Length_{AB}$, Thorax Width = $Length_{CF}$,
2. Right wing lengths: costum = $Length_{CD}$, termen = $Length_{DE}$, dorsum = $Length_{CE}$
3. Left wing lengths: costum = $Length_{FG}$, termen = $Length_{GH}$, dorsum = $Length_{FH}$)

We modeled body-volume as a spindle shaped bicone

$$Body\ Volume(mm^3)=\frac{1}{3}*\Pi\left(\frac{Thorax\ Width}{2}\right)^2(mm^2)*BodyLength(mm)\quad ,$$

and obtained body-mass (hereafter BM) using density of water as follows:

$$BM(gm)=BodyVolume(mm^3)*\left(\frac{1}{1000}\frac{gm}{mm^3}\right)$$

Therefore, body mass and volumes are essentially the same in this study.

Wing area (hereafter WA) was calculated from a triangle defined by the 3 landmarks CDE and FGH.

$$WA(mm^2)=\sqrt{((s*(s-L_{costum})*(s-L_{termen})*(s-L_{dorsum})))}$$

where, $s=\frac{L_{costum}+L_{termen}+L_{dorsum}}{2}$

Wing-loading (hereafter WL) was obtained from

$$WL\left(\frac{gm}{mm^2}\right)=\frac{BM(gm)}{WA(mm^2)}\quad .$$

Wing breadth was obtained from

$$WingBreadth(mm)=\frac{2*WA(mm^2)}{L_{costum}(mm)}$$

Wing aspect-ratio (hereafter AR) was obtained from:

$$AR=\frac{L_{costum}(mm)}{WingBreadth(mm)}\quad .$$

The left and right wing dimensions were used to assess image distortion and subsequently averaged for all analysis (Mungee & Athreya (2019)

### Environmental gradient analysis

We used 4 environmental variables including mean annual temperature (MAT), mean annual precipitation (APPT), productivity (EVI: enhanced vegetation index, and NDVI: normalised vegetation index) and air density (AD). MAT and APPT with a spatial resolution of 1 $km^2$ were downloaded from ***worldclim*** (https://www.worldclim.org/) for the years 2004-2014. EVI and NDVI were obtained from NASA's MODIS satellite products (MOD13Q1) with a resolution of 250 m. Poor pixel quality values were removed before arriving at elevation specific mean values. Air density was calculated for the elevational communities using a global pressure profile and local temperature values using

Air density was calculated as:

$$AD = \frac{P_{Ele} * M}{R * T_{Ele}} \quad ,$$

where $$P_{Ele} = P_{Std} * 1 - \left(\frac{L * Ele}{T_{Std}}\right)^{\left(\frac{gM}{R * T_{Ele}}\right)} \quad ,$$

where $P_{Std}$ = sea level standard atmospheric pressure, 101325 Pa
L = temperature lapse rate, 0.0065 K/m
Ele = elevation (in meters above sea level) for the sampling location
$T_{Std}$ = sea level standard temperature, 288.15 K
g = earth-surface gravitational acceleration, 9.80665 $m/s^2$
M = molar mass of dry air, 0.0289644 kg/mol
R = universal gas constant, 8.31447 J/(mol.K)
$T_{Ele}$ = temperature for the sampling location (K; Obtained from worldclim)

EVI was used over NDVI since the two are strongly correlated and EVI has higher sensitivity at higher biomass.

The four environmental variables were strongly correlated (**Figure 3** below) with all showing a decreasing trend with elevation. The first two axes of a principal component analysis (PCA) explained 91.4 % and 7.7 % of the variance. Therefore, we used PC-1 as a composite environmental variable, or *Environment* (in italics) in all subsequent analysis.

Details on the variables, PC scores and correlations between the variables are provided in **Table 4 and Figure 4** below.

The *Environment* has a strong linear correlation with elevation (**Figure 5** below).

**Table 1. Summary of the hawkmoth sample diversity**

| | |
|---|---|
| Total number of images recorded | 4808 |
| No. of images with species level id (diversity data set) | 4731 |
| No. of species in the diversity data | 80 |
| No. of genera in the diversity data | 30 |
| Moths not on reference grids | 1385 |
| No.. of images processed for calibration | 3346 |
| No. of images removed due to poor dedistortion | 20 |
| No. of individuals removed due to trait outlier flags | 25 |
| Total number of individuals used in all trait analysis (trait data set) | 3301 |
| No. of species in the trait data | 76 |
| No. of genera in the trait data | 30 |

**Table 2. Summary of elevational sampling data for hawkmoths**

The columns headers are: (elevational) Community, Number of sampling nights, Number of individual hawkmoths in diversity data set (DDS), Number in trait data set (TDS), Observed species counts ($S_{DDS}$ and $S_{TDS}$), Rarefied species counts ($R_{DDS}$ and $R_{TDS}$), Fisher's alpha ($F_{DDS}$ and $F_{TDS}$)

| Community | $N_{NIGHT}$ | $N_{DIV}$ | $N_{TRAIT}$ | $S_{DIV}$ | $S_{TRAIT}$ | $R_{DIV}$ | $R_{TRAIT}$ | $F_{DIV}$ | $F_{TRAIT}$ |
|---|---|---|---|---|---|---|---|---|---|
| **E0200** | 14 | 473 | 429 | 43 | 41 | 39 | 26 | 11.5 | 11.2 |
| **E0500** | 5 | 378 | 298 | 39 | 36 | 37 | 25 | 10.9 | 10.7 |
| **E0700** | 8 | 335 | 271 | 33 | 27 | 33 | 21 | 09.1 | 07.5 |
| **E0900** | 5 | 347 | 224 | 45 | 37 | 44 | 29 | 13.8 | 12.6 |
| **E1100** | 11 | 350 | 248 | 48 | 40 | 46 | 29 | 15.1 | 13.5 |
| **E1300** | 9 | 332 | 208 | 31 | 25 | 31 | 20 | 08.4 | 07.4 |
| **E1500** | 8 | 340 | 173 | 40 | 34 | 39 | 28 | 11.8 | 12.7 |
| **E1700** | 4 | 359 | 103 | 36 | 24 | 35 | 24 | 10.0 | 09.8 |
| **E1900** | 7 | 376 | 296 | 40 | 35 | 38 | 24 | 11.3 | 10.3 |
| **E2100** | 8 | 323 | 207 | 29 | 27 | 29 | 21 | 07.7 | 08.3 |
| **E2300** | 10 | 359 | 299 | 27 | 25 | 26 | 17 | 06.8 | 06.5 |
| **E2500** | 6 | 363 | 249 | 23 | 20 | 22 | 14 | 05.5 | 05.1 |
| **E2700** | 12 | 396 | 296 | 32 | 31 | 30 | 20 | 08.2 | 08.7 |

**Table 3. Species trait data summary**

The columns are species mean and standard deviation (SD) of body mass, wing loading (x$10^{-2}$) Aspect Ratio, $N_{DIV}$: number of individuals in diversity data set, $R_{DIV}$: relative abundance in diversity data set, and $N_{TRAIT}$: number of individuals in trait data set.

| | Body mass | | Wing Load (x$10^{-2}$) | | Aspect Ratio | | $N_{DIV}$ | $R_{DIV}$ | $N_{TRAIT}$ |
|---|---|---|---|---|---|---|---|---|---|
| **Taxon** | **Mean** | **SD** | **Mean** | **SD** | **Mean** | **SD** | | | |
| *Macroglossinae_Acosmerycoides_harterti* | 1.20 | 0.25 | 0.52 | 0.09 | 3.90 | 0.34 | 117 | 2.47 | 101 |
| *Macroglossinae_Acosmeryx_anceus* | 1.30 | 0.26 | 0.61 | 0.11 | 3.31 | 0.22 | 20 | 0.42 | 17 |
| *Macroglossinae_Acosmeryx_naga* | 2.56 | 0.49 | 0.63 | 0.10 | 3.45 | 0.26 | 403 | 8.52 | 259 |
| *Macroglossinae_Acosmeryx_omissa* | 1.88 | 0.38 | 0.62 | 0.10 | 3.23 | 0.19 | 237 | 5.01 | 147 |
| *Macroglossinae_Acosmeryx_sericeus* | 2.12 | 0.51 | 0.67 | 0.13 | 3.25 | 0.21 | 113 | 2.39 | 94 |
| *Macroglossinae_Acosmeryx_shervillii* | 1.98 | 0.61 | 0.62 | 0.14 | 3.28 | 0.26 | 229 | 4.84 | 175 |
| *Macroglossinae_Acosmeryx_sinjaevi* | 1.92 | 0.53 | 0.63 | 0.17 | 3.28 | 0.17 | 16 | 0.34 | 11 |
| *Macroglossinae_Ampelophaga_ampelospecies* | NA | NA | NA | NA | NA | NA | 2 | 0.01 | 0 |
| *Macroglossinae_Ampelophaga_dolichoides* | 1.27 | 0.23 | 0.48 | 0.06 | 3.50 | 0.25 | 67 | 1.42 | 57 |
| *Macroglossinae_Ampelophaga_khasiana* | 1.66 | 0.30 | 0.53 | 0.06 | 3.84 | 0.32 | 136 | 2.87 | 99 |
| *Macroglossinae_Ampelophaga_rubiginosa* | 1.58 | 0.38 | 0.58 | 0.15 | 3.43 | 0.25 | 12 | 0.25 | 10 |
| *Macroglossinae_Angonyx_testacea* | 0.77 | 0.09 | 0.54 | 0.01 | 2.88 | 0.13 | 2 | 0.04 | 2 |
| *Macroglossinae_Cechenena_aegrota* | 1.22 | 0.26 | 0.56 | 0.09 | 3.16 | 0.16 | 33 | 0.70 | 30 |
| *Macroglossinae_Cechenena_helops* | 2.51 | 0.45 | 0.79 | 0.11 | 3.44 | 0.58 | 10 | 0.21 | 9 |
| *Macroglossinae_Cechetra_lineosa* | 2.07 | 0.44 | 0.57 | 0.08 | 3.72 | 0.24 | 806 | 17.04 | 523 |
| *Macroglossinae_Cechetra_minor* | 1.46 | 0.22 | 0.60 | 0.11 | 3.51 | 0.23 | 29 | 0.61 | 23 |
| *Macroglossinae_Cechetra_scotti* | 2.45 | 0.53 | 0.61 | 0.09 | 3.83 | 0.22 | 180 | 3.80 | 125 |
| *Macroglossinae_Cechetra_subangustata* | 2.23 | 0.46 | 0.57 | 0.08 | 3.77 | 0.29 | 222 | 4.69 | 179 |
| *Macroglossinae_Daphnis_hypothous* | 3.47 | 1.32 | 0.89 | 0.26 | 3.57 | 0.21 | 11 | 0.23 | 11 |
| *Macroglossinae_Eupanacra_busiris* | 0.76 | NA | 0.76 | NA | 5.75 | NA | 1 | 0.02 | 1 |
| *Macroglossinae_Eupanacra_perfecta* | 0.54 | 0.11 | 0.58 | 0.14 | 3.61 | 0.35 | 17 | 0.36 | 15 |
| *Macroglossinae_Eupanacra_sinuata* | 0.79 | 0.20 | 0.56 | 0.10 | 3.47 | 0.21 | 163 | 3.45 | 112 |
| *Macroglossinae_Eupanacra_variolosa* | 0.66 | NA | 0.70 | NA | 4.20 | NA | 1 | 0.02 | 1 |
| *Macroglossinae_Hippotion_boerhaviae* | 0.57 | 0.08 | 0.53 | 0.09 | 3.67 | 0.19 | 17 | 0.36 | 13 |
| *Macroglossinae_Hippotion_celerio* | 0.86 | 0.09 | 0.63 | 0.05 | 3.69 | 0.19 | 6 | 0.13 | 3 |
| *Macroglossinae_Macroglossum_macrosp1* | NA | NA | NA | NA | NA | NA | 1 | 0.01 | 0 |
| *Macroglossinae_Macroglossum_macrosp2* | 1.26 | 0.35 | 1.01 | 0.16 | 3.28 | 0.31 | 3 | 0.06 | 2 |
| *Macroglossinae_Nephele_hespera* | 1.29 | 0.36 | 0.71 | 0.21 | 3.09 | 0.15 | 18 | 0.38 | 17 |
| *Macroglossinae_Pergesa_acteus* | 1.21 | 0.23 | 0.75 | 0.16 | 3.51 | 0.21 | 13 | 0.27 | 12 |
| *Macroglossinae_Rhagastis_acuta* | 0.79 | 0.16 | 0.55 | 0.11 | 3.02 | 0.23 | 11 | 0.23 | 10 |
| *Macroglossinae_Rhagastis_albomarginatus* | 1.32 | 0.32 | 0.48 | 0.07 | 3.38 | 0.12 | 38 | 0.80 | 16 |
| *Macroglossinae_Rhagastis_castor* | 1.11 | 0.31 | 0.49 | 0.09 | 3.19 | 0.21 | 57 | 1.20 | 39 |
| *Macroglossinae_Rhagastis_confusa* | 1.06 | 0.25 | 0.45 | 0.07 | 3.32 | 0.27 | 102 | 2.16 | 64 |
| *Macroglossinae_Rhagastis_gloriosa* | 1.21 | 0.26 | 0.46 | 0.07 | 3.02 | 0.30 | 64 | 1.35 | 54 |
| *Macroglossinae_Rhagastis_lunata* | 1.51 | 0.35 | 0.56 | 0.10 | 3.07 | 0.17 | 111 | 2.35 | 88 |
| *Macroglossinae_Rhagastis_olivacea* | 1.01 | 0.18 | 0.48 | 0.07 | 3.17 | 0.28 | 66 | 1.40 | 39 |
| *Macroglossinae_Rhagastis_velata* | 0.95 | 0.25 | 0.59 | 0.12 | 2.95 | 0.14 | 5 | 0.11 | 3 |
| *Macroglossinae_Theretra_alecto* | 1.72 | 0.42 | 0.66 | 0.17 | 3.35 | 0.18 | 20 | 0.42 | 14 |
| *Macroglossinae_Theretra_boisduvalii* | 2.21 | 0.46 | 0.67 | 0.12 | 3.56 | 0.23 | 226 | 4.78 | 174 |
| *Macroglossinae_Theretra_clotho* | 2.01 | 0.43 | 0.79 | 0.18 | 3.60 | 0.17 | 179 | 3.78 | 140 |
| *Macroglossinae_Theretra_griseomarginata* | 0.50 | 0.05 | 0.44 | 0.03 | 2.88 | 0.22 | 4 | 0.08 | 2 |
| *Macroglossinae_Theretra_latreillii* | 1.15 | 0.25 | 0.58 | 0.12 | 3.23 | 0.18 | 5 | 0.11 | 5 |
| *Macroglossinae_Theretra_nessus* | 2.99 | 0.57 | 0.69 | 0.12 | 3.57 | 0.19 | 40 | 0.85 | 33 |
| *Macroglossinae_Theretra_oldenlandiae* | 0.79 | 0.16 | 0.50 | 0.06 | 3.54 | 0.17 | 19 | 0.40 | 15 |
| *Macroglossinae_Theretra_pallicosta* | 1.49 | 0.35 | 0.66 | 0.14 | 3.43 | 0.16 | 8 | 0.17 | 5 |
| *Macroglossinae_Theretra_suffusa* | 1.45 | NA | 0.70 | NA | 3.62 | NA | 1 | 0.02 | 1 |
| *Smerinthinae_Ambulyx_liturata* | 2.47 | 0.51 | 0.48 | 0.07 | 3.93 | 0.40 | 123 | 2.60 | 64 |
| *Smerinthinae_Ambulyx_maculifera* | 1.66 | 0.41 | 0.46 | 0.11 | 3.60 | 0.17 | 20 | 0.42 | 12 |
| *Smerinthinae_Ambulyx_moorei* | 1.48 | 0.24 | 0.41 | 0.02 | 3.57 | 0.13 | 2 | 0.04 | 2 |

| | | | | | | | | | |
|---|---|---|---|---|---|---|---|---|---|
| *Smerinthinae_Ambulyx_ochracea* | 1.13 | 0.26 | 0.38 | 0.07 | 3.62 | 0.24 | 101 | 2.13 | 73 |
| *Smerinthinae_Ambulyx_pseudoclavata* | 1.92 | 0.52 | 0.42 | 0.11 | 3.52 | 0.27 | 10 | 0.21 | 9 |
| *Smerinthinae_Ambulyx_sericeipennis* | 2.22 | 0.15 | 0.39 | 0.02 | 3.59 | 0.20 | 5 | 0.11 | 3 |
| *Smerinthinae_Ambulyx_substrigilis* | 1.73 | 0.34 | 0.43 | 0.05 | 3.43 | 0.19 | 48 | 1.01 | 28 |
| *Smerinthinae_Ambulyx_tobii* | 2.24 | 0.36 | 0.46 | 0.04 | 3.55 | 0.16 | 34 | 0.72 | 21 |
| *Smerinthinae_Amplypterus_mansoni* | 4.19 | 0.53 | 0.63 | 0.07 | 3.58 | 0.18 | 33 | 0.70 | 13 |
| *Smerinthinae_Amplypterus_panopus* | 3.22 | 0.71 | 0.53 | 0.01 | 3.48 | 0.01 | 3 | 0.06 | 2 |
| *Smerinthinae_Callambulyx_junonia* | 1.56 | 0.29 | 0.44 | 0.05 | 3.26 | 0.20 | 7 | 0.15 | 4 |
| *Smerinthinae_Callambulyx_poecilus* | 0.88 | 0.22 | 0.45 | 0.09 | 3.71 | 0.32 | 14 | 0.30 | 10 |
| *Smerinthinae_Callambulyx_rubricosa* | 2.75 | 0.77 | 0.56 | 0.13 | 3.64 | 0.42 | 46 | 0.97 | 26 |
| *Smerinthinae_Clanis_schwartzi* | NA | NA | NA | NA | NA | NA | 1 | 0.01 | 0 |
| *Smerinthinae_Clanis_titan* | NA | NA | NA | NA | NA | NA | 1 | 0.01 | 0 |
| *Smerinthinae_Clanis_undulosa* | 4.87 | 0.93 | 0.81 | 0.17 | 5.06 | 0.79 | 30 | 0.63 | 13 |
| *Smerinthinae_Craspedortha_porphyria* | 0.51 | 0.13 | 0.63 | 0.32 | 3.25 | 1.09 | 21 | 0.44 | 15 |
| *Smerinthinae_Cypa_decolor* | 0.75 | NA | 0.68 | NA | 3.25 | NA | 1 | 0.02 | 1 |
| *Smerinthinae_Daphnusa_sinocontinentalis* | 1.13 | 0.24 | 0.43 | 0.10 | 2.79 | 0.17 | 7 | 0.15 | 6 |
| *Smerinthinae_Dolbina_inexacta* | 1.28 | 0.19 | 0.58 | 0.07 | 3.36 | 0.20 | 14 | 0.30 | 11 |
| *Smerinthinae_Marumba_cristata* | 2.56 | 0.72 | 0.51 | 0.11 | 3.35 | 0.34 | 15 | 0.32 | 9 |
| *Smerinthinae_Marumba_dyras* | 2.19 | 0.46 | 0.56 | 0.11 | 3.23 | 0.31 | 34 | 0.72 | 30 |
| *Smerinthinae_Marumba_echephron* | 2.32 | 0.81 | 0.62 | 0.19 | 3.85 | 0.81 | 11 | 0.23 | 4 |
| *Smerinthinae_Marumba_spectabilis* | 2.10 | 0.08 | 0.73 | 0.15 | 3.48 | 0.99 | 2 | 0.04 | 2 |
| *Smerinthinae_Rhodoprasina_rhodospecies* | 1.50 | 0.23 | 0.53 | 0.07 | 3.81 | 0.28 | 9 | 0.19 | 5 |
| *Sphinginae_Acherontia_lachesis* | 4.47 | 1.16 | 1.12 | 0.22 | 3.98 | 0.62 | 41 | 0.87 | 26 |
| *Sphinginae_Acherontia_styx* | 2.49 | 0.19 | 0.84 | 0.38 | 3.95 | 0.73 | 3 | 0.06 | 2 |
| *Sphinginae_Agrius_convolvuli* | 2.06 | 0.63 | 0.88 | 0.25 | 4.36 | 0.89 | 60 | 1.27 | 14 |
| *Sphinginae_Apocalypsis_velox* | 3.76 | 0.99 | 0.63 | 0.14 | 3.58 | 0.21 | 35 | 0.74 | 29 |
| *Sphinginae_Megacorma_obliqua* | 4.25 | 0.30 | 0.92 | 0.05 | 3.35 | 0.31 | 3 | 0.06 | 3 |
| *Sphinginae_Meganoton_analis* | 4.08 | 0.71 | 0.57 | 0.09 | 3.44 | 0.22 | 66 | 1.40 | 47 |
| *Sphinginae_Meganoton_rubescens* | 2.75 | 0.77 | 0.50 | 0.09 | 3.20 | 0.21 | 28 | 0.59 | 20 |
| *Sphinginae_Psilogramma_increta* | 3.39 | 0.55 | 0.64 | 0.09 | 3.63 | 0.11 | 7 | 0.15 | 6 |
| *Sphinginae_Psilogramma_menephron* | 3.16 | 0.88 | 0.63 | 0.09 | 3.49 | 0.44 | 55 | 1.16 | 41 |

**Table 4. Environmental variables of sampled communities**

The columns are Plot: elevational community, (actual) Elevation, MAT: mean annual temperature, APPT: mean annual precipitation, EVI: enhanced vegetation index (productivity), NDVI : normalised difference vegetation index (productivity), AD: air density, PC1: principal component 1 and PC2. PC1 was used as the environmental variable.

| Plot | Elevation | MAT | APPT | EVI | *NDVI* * | AD | Score on PC1 | Score on PC2 |
|---|---|---|---|---|---|---|---|---|
| **E0200** | 268 | 23.6 | 2308 | 0.4373 | *0.7024* | 1.152 | -2.88 | -0.21 |
| **E0500** | 502 | 22.3 | 2283 | 0.3812 | *0.6977* | 1.125 | -2.03 | 0.52 |
| **E0700** | 715 | 21.3 | 2268 | 0.3881 | *0.6471* | 1.101 | -1.80 | 0.23 |
| **E0900** | 897 | 19.8 | 2185 | 0.3555 | *0.6280* | 1.082 | -1.11 | 0.57 |
| **E1100** | 1106 | 19.8 | 2185 | 0.3599 | *0.6494* | 1.055 | -0.99 | 0.36 |
| **E1300** | 1305 | 19.4 | 2154 | 0.4026 | *0.6218* | 1.031 | -1.12 | -0.50 |
| **E1500** | 1503 | 16.8 | 1849 | 0.3207 | *0.5607* | 1.016 | 0.38 | 0.54 |
| **E1700** | 1712 | 16.8 | 1849 | 0.3597 | *0.5958* | 0.990 | 0.20 | -0.24 |
| **E1900** | 1902 | 16.1 | 1754 | 0.4078 | *0.6692* | 0.969 | 0.11 | -1.21 |
| **E2100** | 2103 | 14.4 | 1526 | 0.3509 | *0.5807* | 0.951 | 1.21 | -0.53 |
| **E2300** | 2318 | 13.4 | 1394 | 0.2981 | *0.5398* | 0.929 | 2.10 | 0.14 |
| **E2500** | 2499 | 12.4 | 1274 | 0.3142 | *0.6715* | 0.911 | 2.33 | -0.31 |
| **E2700** | 2770 | 10.3 | 1042 | 0.2397 | *0.4878* | 0.895 | 3.63 | 0.65 |

**Figure 1. Nightly hawkmoth visitation rates at light screens.**
The X-axis is sequence number of the night at that location, and the Y-axis represents the number of hawkmoths. The blue shaded region represents the number of individuals of the most abundant species on that day.

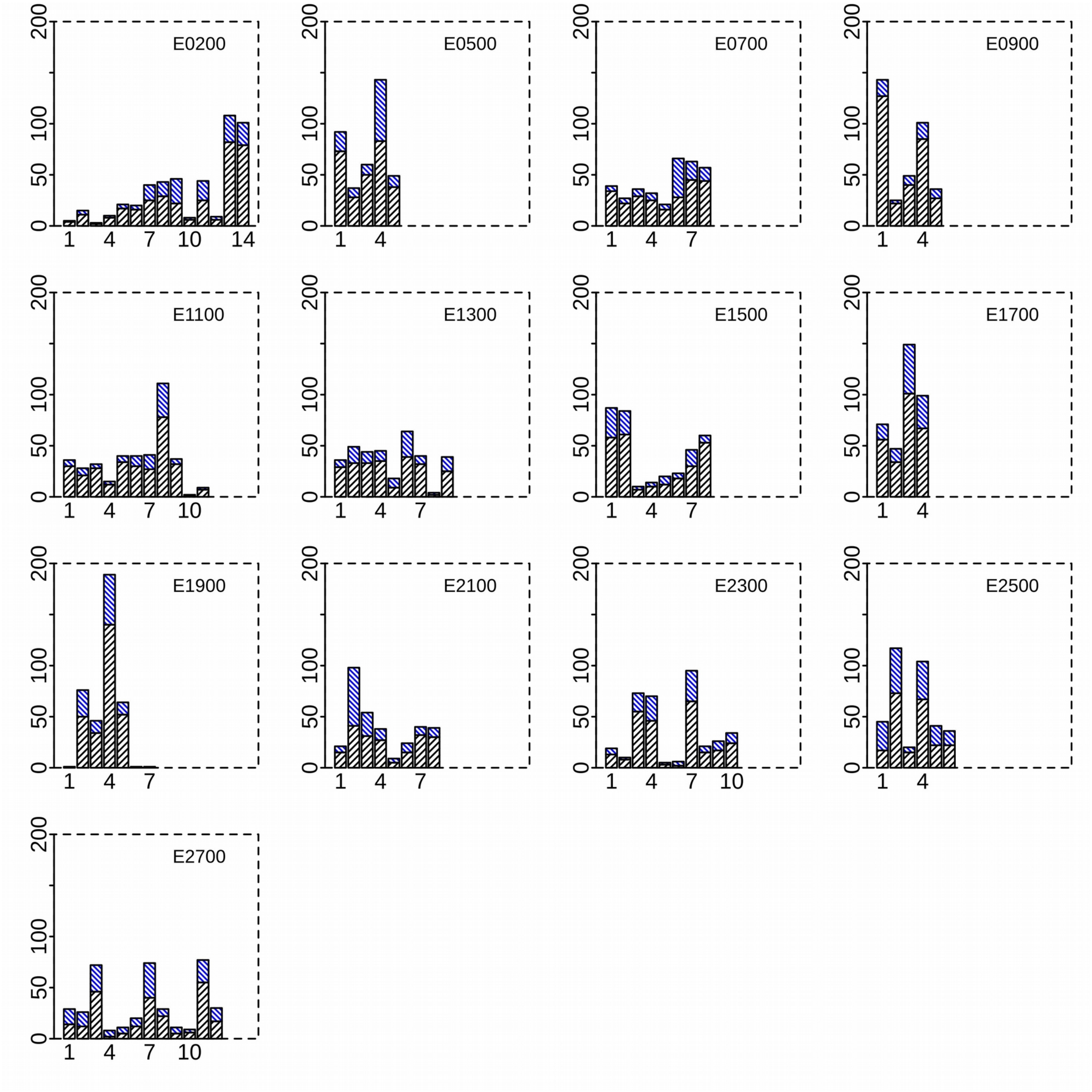

**Figure 2. Trait data images of hawkmoths**

Individuals were imaged against a reference background grid used as UV illuminated light screen. The rectangular grids were used to calibrate the image for shape and size. The 8 landmarks A-H were used to measure the body and wing dimensions.

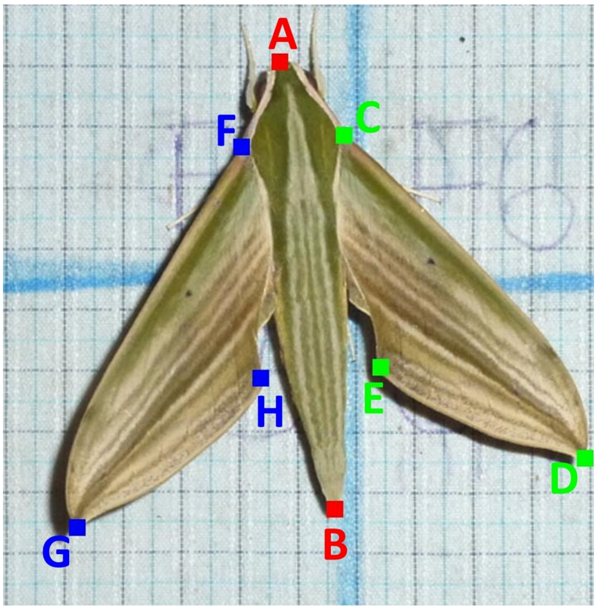

**Figure 3. Relationships between the environmental variables.**

The environmental variables are EVI: enhanced vegetation index, NDVI: normalized difference vegetation index, MAT: mean annual temperature, APPT, mean annual precipitation, and air density. Linear regression coefficients between some of the variables are listed in the table below.

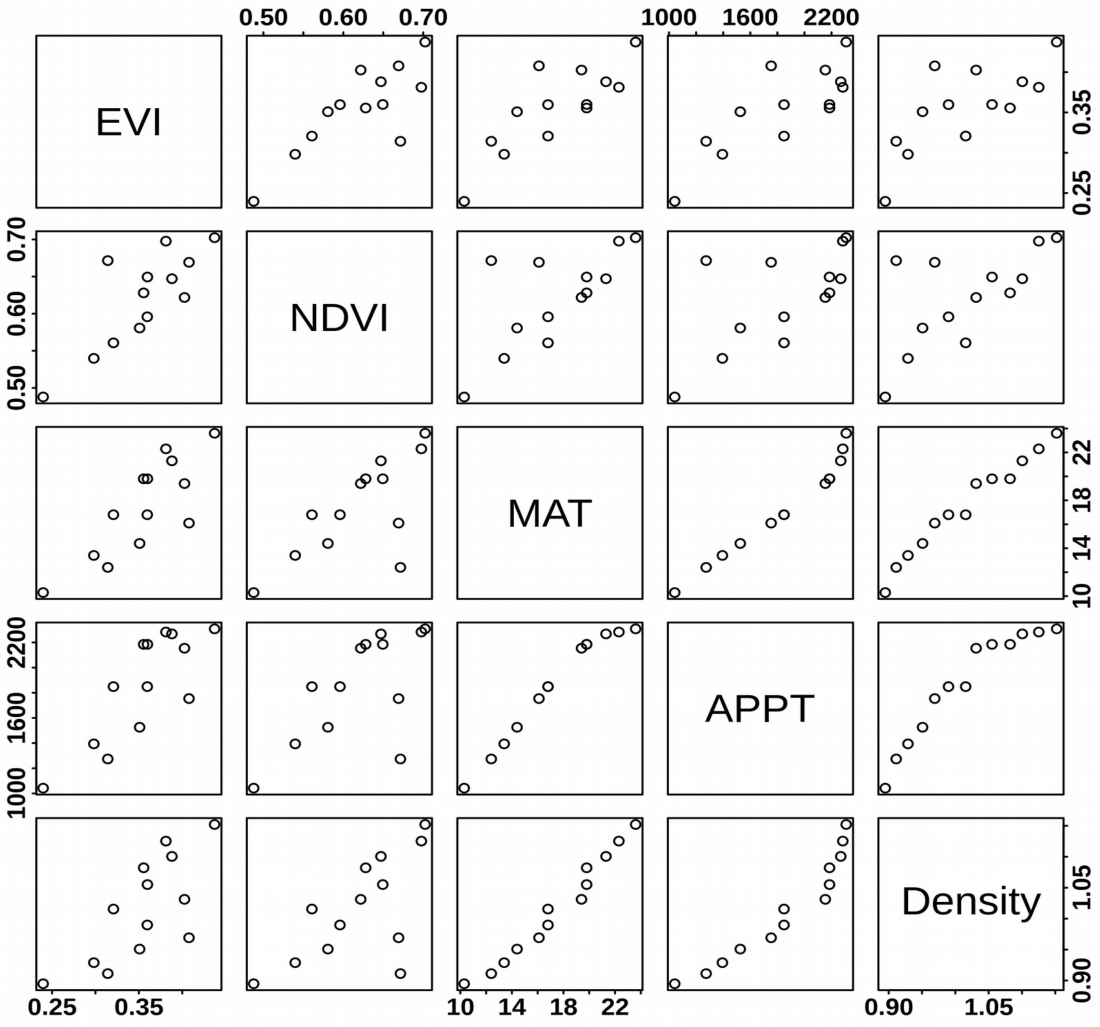


| Model | Slope | Intercept | R.squared | p.value |
|---|---|---|---|---|
| EVI ~ Elevation | $-5.04 \times 10^{-5}$ | $4.31 \times 10^{-1}$ | 0.53 | < 0.005 |
| NDVI ~ Elevation | $-5.42 \times 10^{-5}$ | $7.01 \times 10^{-1}$ | 0.39 | < 0.05 |
| MAT ~ Elevation | $-5.10 \times 10^{-3}$ | $2.51 \times 10^{1}$ | 0.98 | < 0.005 |
| APPT ~ Elevation | $-5.25 \times 10^{-1}$ | $2.64 \times 10^{3}$ | 0.92 | < 0.005 |
| AD ~ Elevation | $-1.1 \times 10^{-4}$ | $0.12 \times 10^{1}$ | 0.99 | < 0.005 |
| EVI ~ NDVI | $0.98 \times 10^{-1}$ | $2.72 \times 10^{-1}$ | 0.61 | < 0.005 |
| EVI ~ APPT | $0.66 \times 10^{4}$ | $-4.91 \times 10^{2}$ | 0.62 | < 0.005 |
| EVI ~ AD | $4.6 \times 10^{-1}$ | $-1.12 \times 10^{-1}$ | 0.50 | < 0.005 |

**Figure 4. Principal component analysis of environmental variables**
Four environmental variable were used: AD Air density, APPT mean annual precipitation, EVI Enhanced vegetation index (productivity), MAT Mean Annual Temperature. Axis 1 explained 91.4% of the variation and was used as the environmental gradient in the analysis. . The elevational communities are marked on the plot (see **Table 4** above).

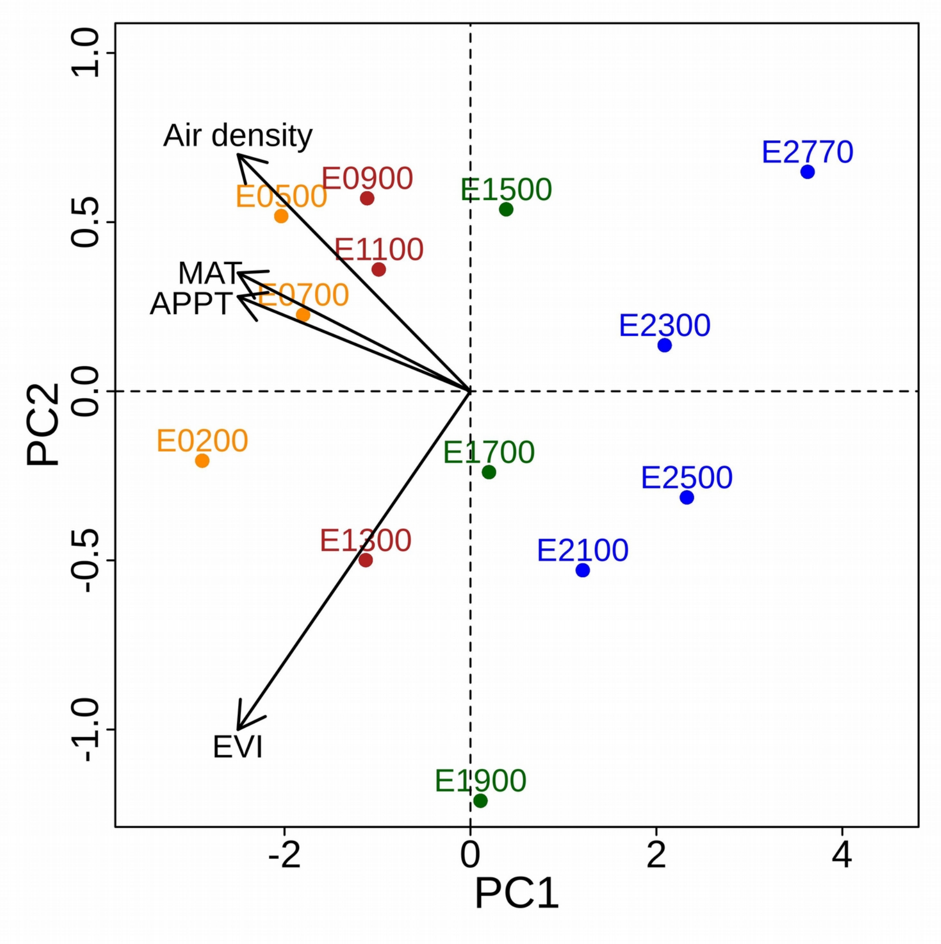


| | **PC1** | **PC2** | **PC3** | **PC4** |
|---|---|---|---|---|
| **EVI** | -0.459 | -0.869 | -0.146 | -0.113 |
| **MAT** | -0.520 | 0.186 | -0.115 | 0.825 |
| **APPT** | -0.514 | 0.169 | 0.803 | -0.250 |
| **AD** | -0.505 | 0.425 | -0.566 | -0.493 |

**Figure 5. Relationship between the composite environmental variable and elevation**
The composite environmental variable is the first principal component, which includes 91.4% of the total variance. The slope and the $R^2$ of the Ordinary least square regression is listed on the plot.

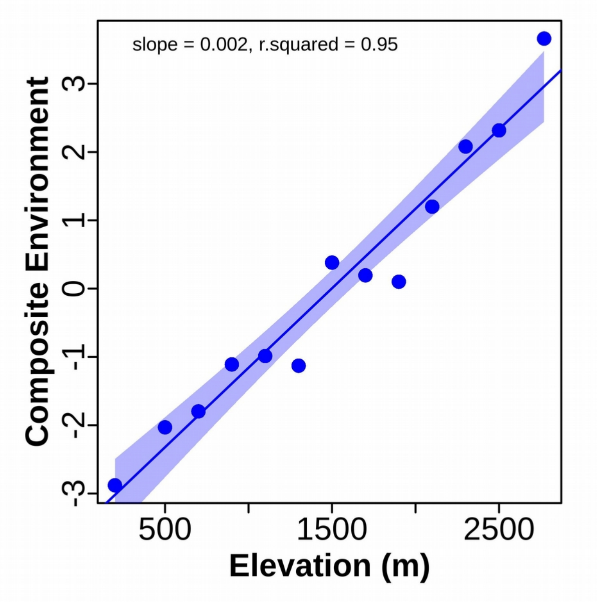

**Figure 6. Hawkmoth taxonomic diversity rarefaction curves.**
Each color represents a different elevational community

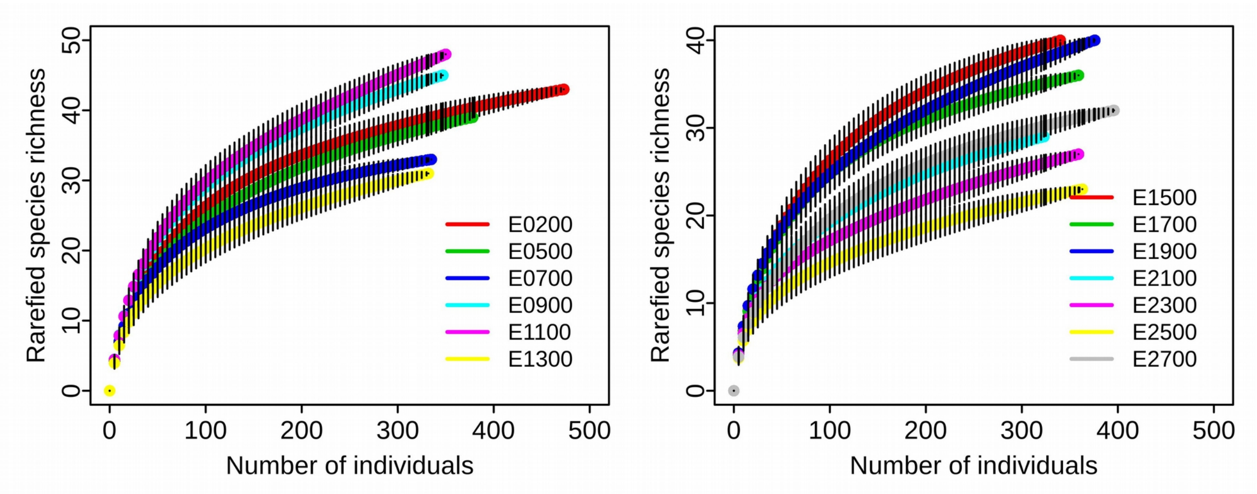

**Figure 7. Hawkmoth functional diversity rarefaction curves.**
Each plot represents a different elevational community.
It can be seen that the overall trait volume is quickly achieved by sample size 50-100. The rest of the data essentially fill in the “intraspecific variability” space.The traits were normalized using community-specific mean and standard deviations prior to analysis. The Rarefactions were performed using function *Rarefaction* from package *evolqg* (Melo et al. 2015, R core development Team, 2013).

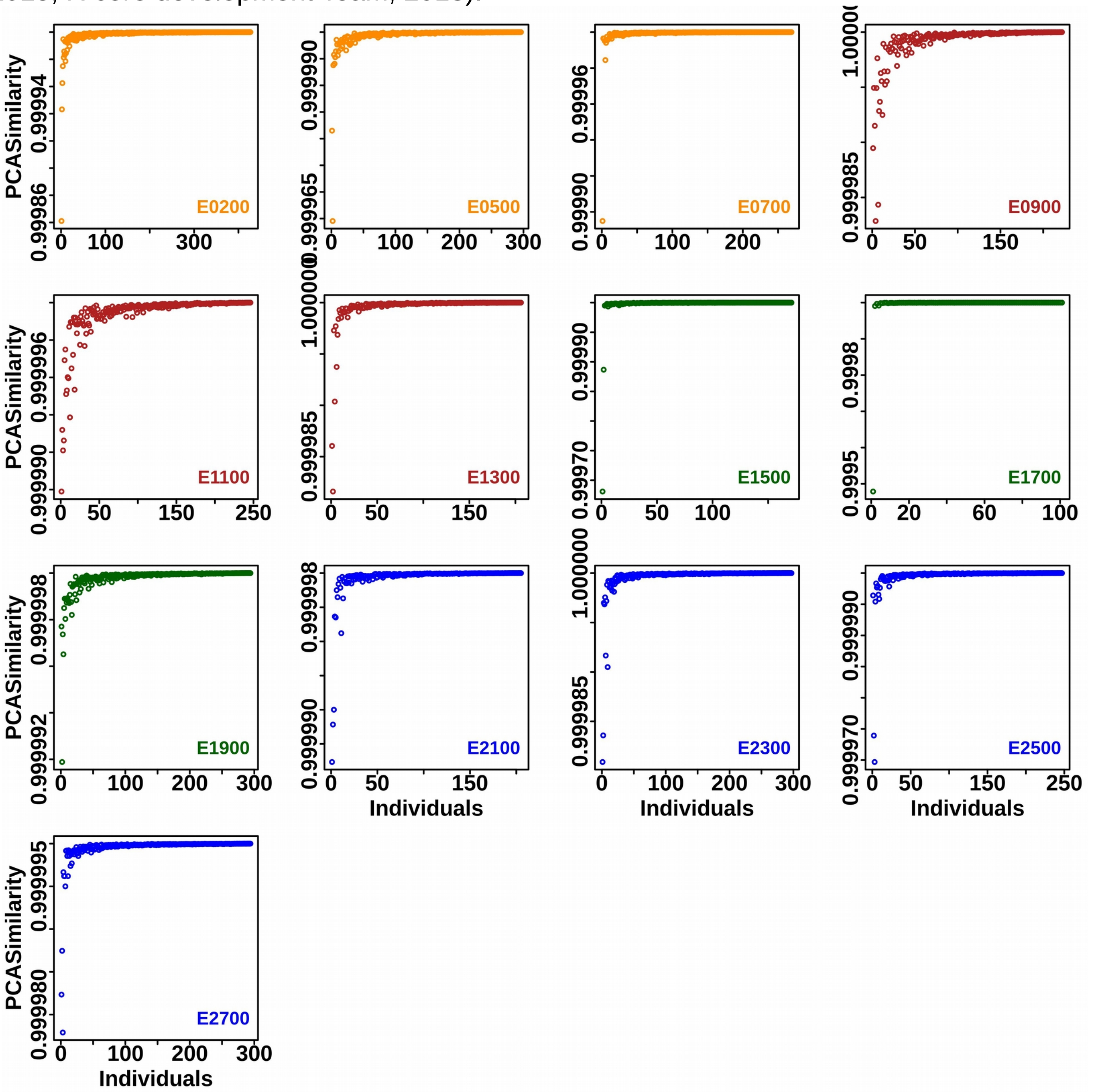
