## Supplementary_02 for "Intraspecific trait variability and community assembly in hawkmoths (Lepidoptera: *Sphingidae*) across an elevational gradient in the eastern Himalayas, India"

We repeated the three major analyses with modified data sets to understand the impact of incomplete data.

**A. diversity dataset with resampling of missing traits from others from the same elevational community**

We could measure traits for only 3301 individuals (the trait data set) out of the total diversity data set of 4731 individuals. Therefore, we redid the analysis by filling in the missing trait data by randomly resampling the traits from others of the same species in the same community

a) Trait variation with *Environment*:
   Figure 1 below; to be compared with Main, Figure 2
b) Trait overlap with *Environment*al distance:
   Figure 2 below; to be compared with Main, Figure 3
c) T-statistics along environmental gradient:
   Figure 3 below; to be compared with Main, Figure 5

**B. trait data set without E1700 data**

E1700 data suffered a disproportionate loss of trait data due to adverse weather conditions during which the moths could not be photographed against the reference grid

a) Trait variation with *Environment*:
   Figure 4 below; to be compared with Main, Figure 2
b) Trait overlap with *Environment*al distance:
   Figure 5 below; to be compared with Main, Figure 3
c) T-statistics along environmental gradient:
   Figure 6 below; to be compared with Main, Figure 5

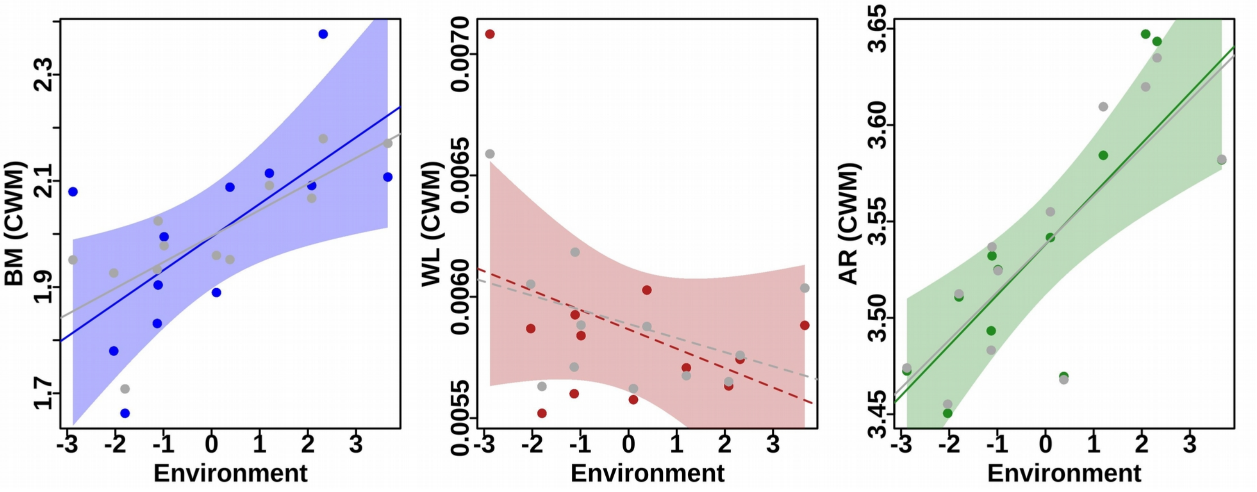


| | Intercept | SE_Intercept | Slope | SE_Slope | Rsquared | p.value |
|---|---|---|---|---|---|---|
| **BM(CWM1) ~ Gradient** | 2.00 | 0.03 | 0.05 | 0.02 | 0.40 | < 0.05* |
| **BM(CWM2) ~ Gradient** | 2.00 | 0.05 | 0.06 | 0.03 | 0.31 | < 0.05* |
| **WL(CWM1) ~ Gradient** | $6 \times 10^{-3}$ | $8 \times 10^{-5}$ | $-4 \times 10^{-5}$ | $4 \times 10^{-5}$ | -0.01 | 0.33 |
| **WL(CWM2) ~ Gradient** | $6 \times 10^{-3}$ | $1 \times 10^{-4}$ | $-7 \times 10^{-5}$ | $6 \times 10^{-5}$ | 0.03 | 0.28 |
| **AR(CWM1) ~ Gradient** | 3.53 | 0.01 | 0.03 | 0.01 | 0.68 | < 0.005** |
| **AR(CWM2) ~ Gradient** | 3.53 | 0.01 | 0.03 | 0.01 | 0.48 | < 0.005** |

**Figure 1. Variation of hawkmoth community traits with *Environment*.**
The plots show the change in community weighted means (CWM) of body mass (BM), wing loading (WL) and wing aspect ratio (AR) plotted against composite E*nvironment* (which has a strong positive correlation with elevation). For each trait, the colored representations are for CWM1 (population mean trait values weighted by local abundance) while the grey are for CWM2 (regional species mean trait values weighted by local abundance). The lines represent the best fits, with the ones significant at 95% shown in solid and the others with dashes.

Using diversity data set with the missing trait values obtained by resampling other individuals of the same species from the same community

To be compared with Main Figure 2

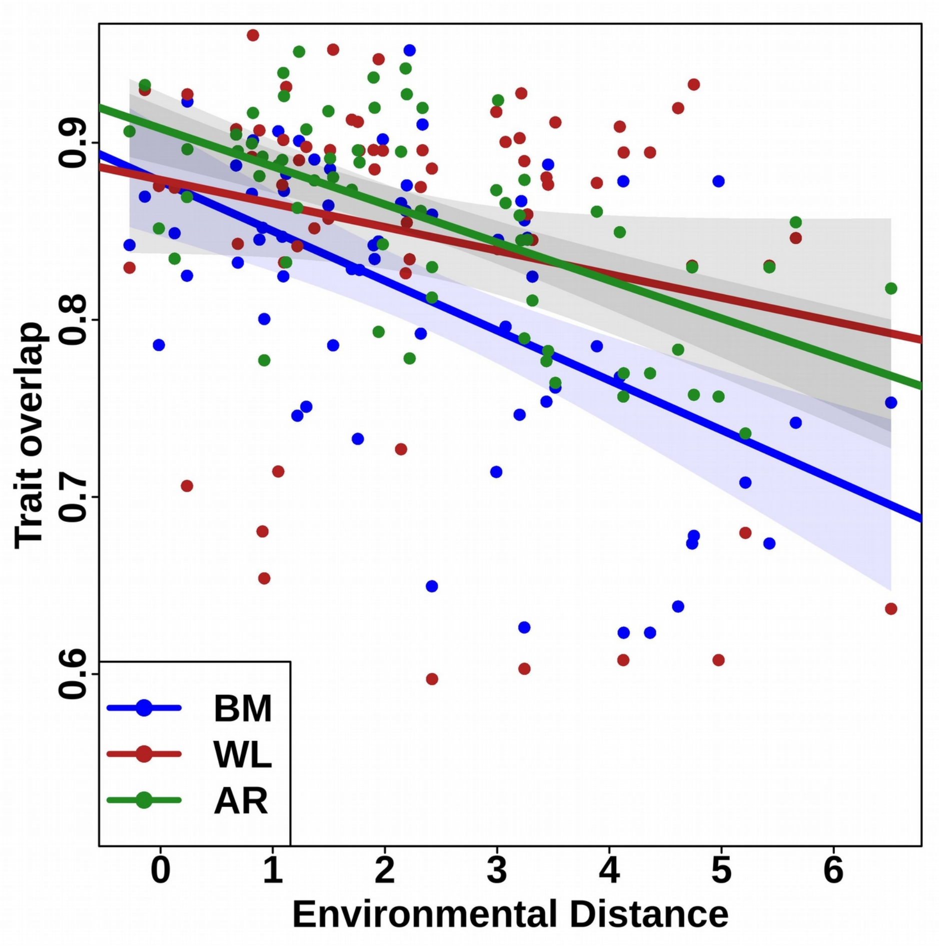


| | Intercept | SE_Intercept | Slope | SE_Slope | Rsquared | p.value |
|---|---|---|---|---|---|---|
| **BM ~ Distance** | 0.88 | 0.02 | $-7.4 \times 10^{-5}$ | $1.4 \times 10^{-5}$ | 0.25 | < 0.005** |
| **WL ~ Distance** | 0.88 | 0.02 | -4.7 x 10-$^{5}$ | $1.6 \times 10^{-5}$ | 0.09 | < 0.005** |
| **AR ~ Distance** | 0.88 | 0.01 | $-5.1 \times 10^{-5}$ | $0.9 \times 10^{-5}$ | 0.27 | < 0.005** |

**Figure 2. Relationship between trait overlap and environmental distance**
The plot shows the scatter and the regression line for the relationship between the overlap in trait distribution functions for pairs of communities and the *Environment*al distance between them. The shaded areas are the 95% errors on the slope. The three traits analysed are body mass (BM), wing loading (WL), and wing aspect ratio (AR). The statistically significant correlation suggests that these are indeed functional traits.

Using diversity data set with the missing trait values obtained by resampling other individuals of the same species from the same community

To be compared with Main Figure 3

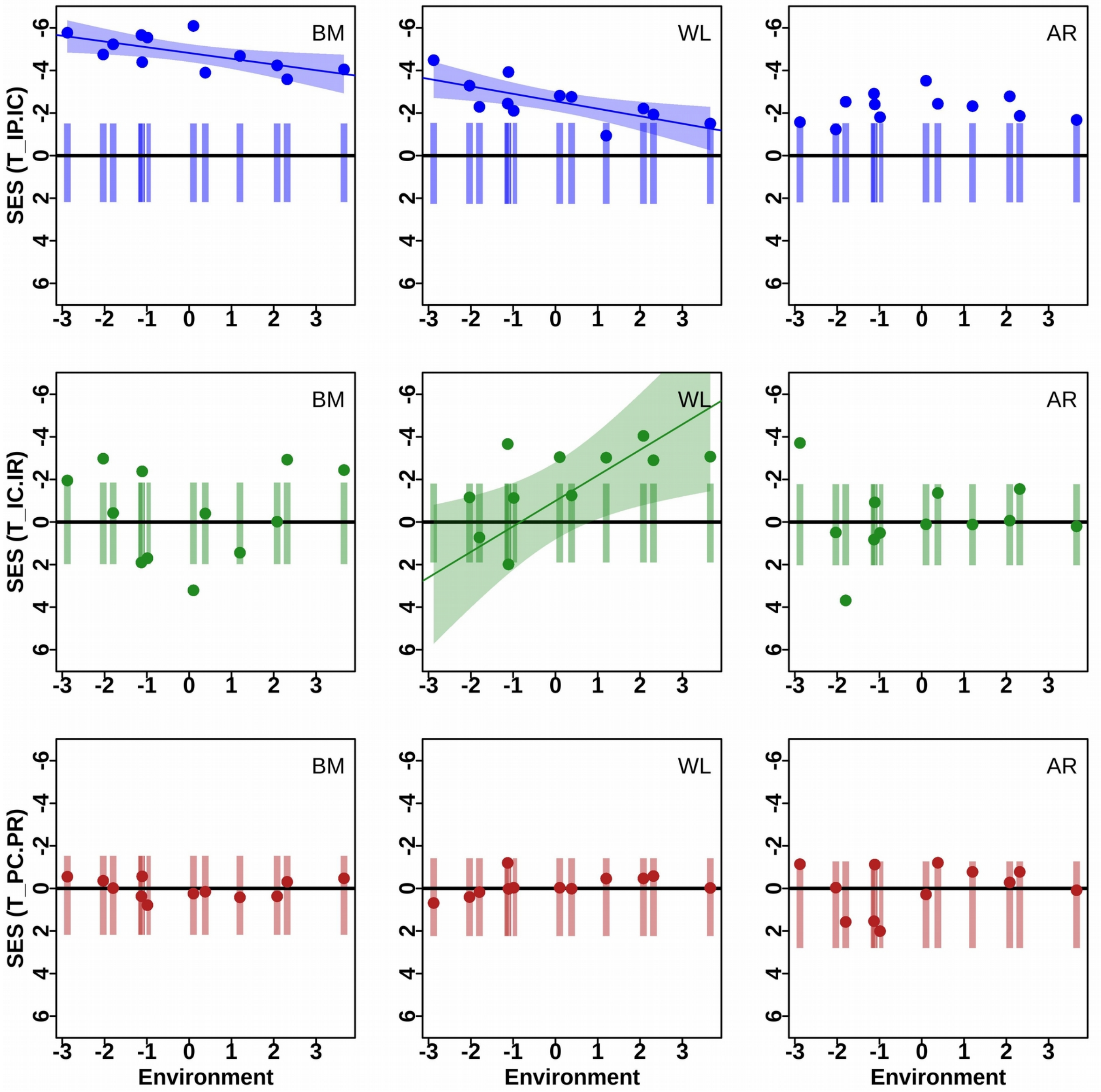


**Figure 3. T-statistics of hawkmoth functional traits across an *Environmental* gradient**

The plots show the T-statistics for 3 traits: body mass (BM), wing loading (WL) and wing aspect ratio (AR), and for each of the 13 elevational communities (along the x-axis) in terms of the standardized effect size (SES) along the y-axis. The vertical bars are the 95% distribution from 999 simulated null communities, and the dots are the observed community values. The three T-statistics variance ratios are (a) $T_{IP/IC}$ — within-population to within-community; (b) $T_{IC/IR}$ — within-community to within-regional, assessed using individual values; and (c) $T_{PC/PR}$ — within-community-wide to within-region, assessed using population-means. Additionally, linear regression fits including errors (shaded region) have been shown for metrics which are significantly correlated (CI > 95%) with the composite *Environmenta*l variable (which is positively correlated with elevation).

Using diversity data set with the missing trait values obtained by resampling other individuals of the same species from the same community

To be compared with Main Figure 5

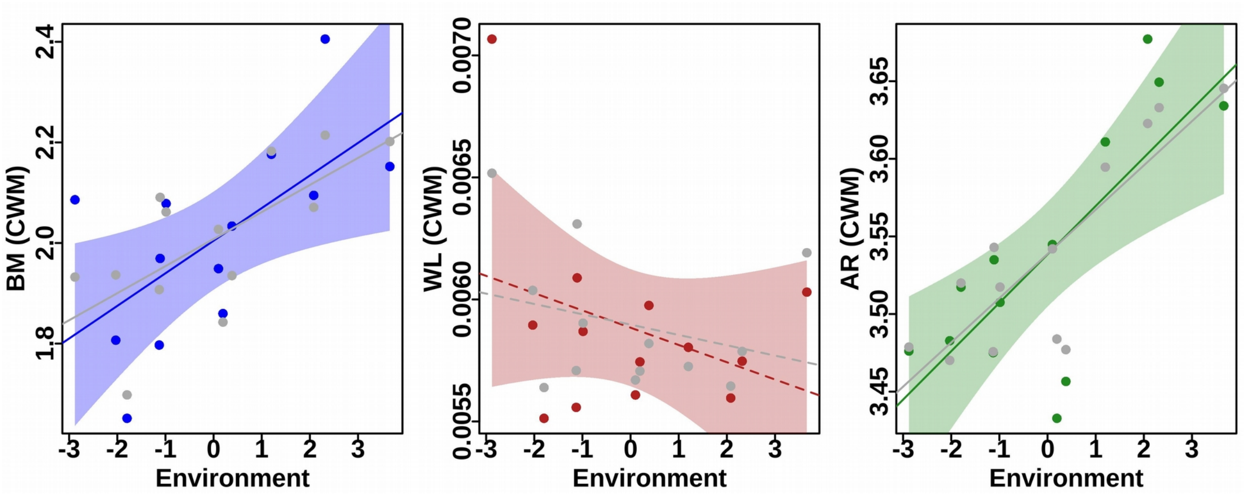


| | Intercept | SE_Intercept | Slope | SE_Slope | Rsquared | p.value |
|---|---|---|---|---|---|---|
| **BM(CWM1) ~ Gradient** | 2.00 | 0.03 | 0.05 | 0.02 | 0.41 | < 0.05 |
| **BM(CWM2) ~ Gradient** | 2.00 | 0.05 | 0.06 | 0.03 | 0.32 | < 0.05 |
| **WL(CWM1) ~ Gradient** | $6 \times 10^{-3}$ | $8 \times 10^{-5}$ | $-4 \times 10^{-5}$ | $4 \times 10^{-5}$ | 0.001 | 0.33 |
| **WL(CWM2) ~ Gradient** | $6 \times 10^{-3}$ | $1 \times 10^{-4}$ | $-7 \times 10^{-5}$ | $6 \times 10^{-5}$ | 0.03 | 0.28 |
| **AR(CWM1) ~ Gradient** | 3.54 | 0.01 | 0.03 | 0.01 | 0.73 | < 0.005 |
| **AR(CWM2) ~ Gradient** | 3.54 | 0.01 | 0.03 | 0.01 | 0.59 | < 0.005 |

**Figure 4: Variation of hawkmoth community traits along an *Environment*.**
The plots show the change in community weighted means (CWM) of body mass (BM), wing loading (WL) and wing aspect ratio (AR) plotted against composite E*nvironment* (which has a strong positive correlation with elevation). For each trait, the colored representations are for CWM1 (population mean trait values weighted by local abundance) while the grey are for CWM2 (regional species mean trait values weighted by local abundance). The lines represent the best fits, with the ones significant at 95% shown in solid and the others with dashes.

Using trait data set without E1700 data

To be compared with Main Figure 2

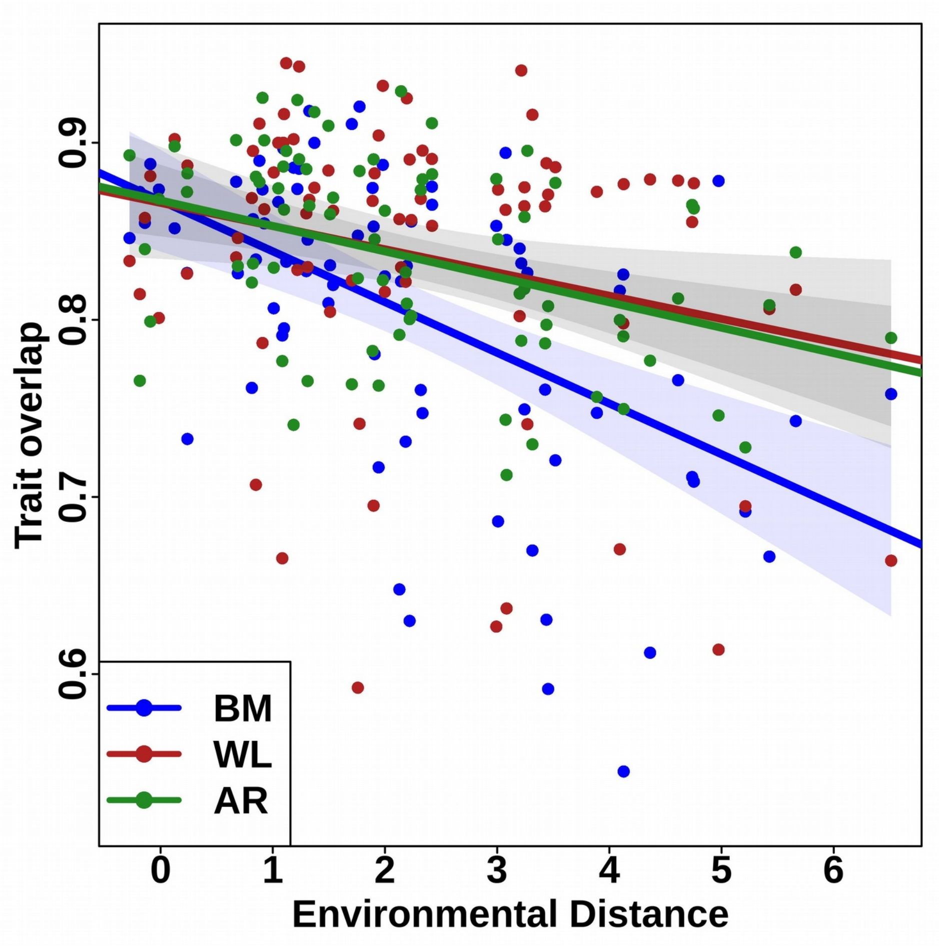


| | Intercept | SE_Intercept | Slope | SE_Slope | Rsquared | p.value |
|---|---|---|---|---|---|---|
| BM ~ Gradient | 0.87 | 0.02 | -0.03 | 0.005 | 0.26 | < 0.005 |
| WL ~ Gradient | 0.87 | 0.02 | -0.01 | 0.006 | 0.05 | < 0.05 |
| AR ~ Gradient | 0.87 | 0.01 | -0.01 | 0.004 | 0.15 | < 0.005 |

**Figure 5: Relationship between trait overlap and environmental distance**
The plot shows the scatter and the regression line for the relationship between the overlap in trait distribution functions for pairs of communities and the *Environmenta*l distance between them. The shaded areas are the 95% errors on the slope. The three traits analysed are body mass (BM), wing loading (WL), and wing aspect ratio (AR). The statistically significant correlation suggests that these are indeed functional traits.

Using trait data set without E1700 data

To be compared with Main Figure 3

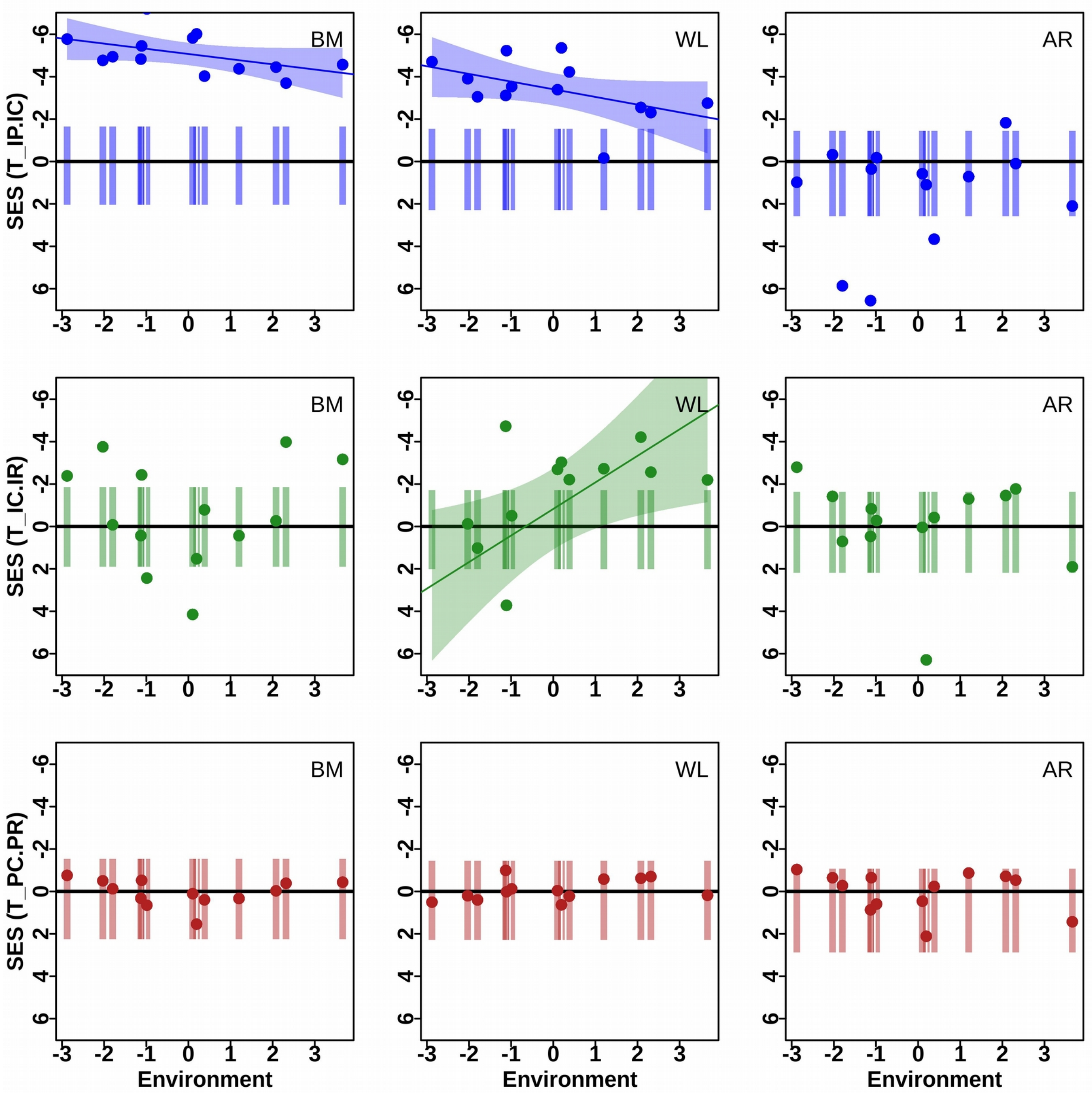


**Figure 6. T-statistics of hawkmoth functional traits across an *Environmental* gradient**

The plots show the T-statistics for 3 traits: body mass (BM), wing loading (WL) and wing aspect ratio (AR), and for each of the 13 elevational communities (along the x-axis) in terms of the standardized effect size (SES) along the y-axis. The vertical bars are the 95% distribution from 999 simulated null communities, and the dots are the observed community values. The three T-statistics variance ratios are (a) $T_{IP/IC}$ — within-population to within-community; (b) $T_{IC/IR}$ — within-community to within-regional, assessed using individual values; and (c) $T_{PC/PR}$ — within-community-wide to within-region, assessed using population-means. Additionally, linear regression fits including errors (shaded region) have been shown for metrics which are significantly correlated (CI > 95%) with the composite *Environment*al variable (which is positively correlated with elevation).

Using trait data set without E1700 data

To be compared with Main Figure 5
