## Supplementary_03 for "Intraspecific trait variability and community assembly in hawkmoths (Lepidoptera: *Sphingidae*) across an elevational gradient in the eastern Himalayas, India"

**Table 1** Null models used to calculate the statistical significance for T-statistics (adapted from Taudière & Violle 2015).

| | **Null model** | **Null hypothesis** | **Randomization** | **Alternate hypothesis** |
|---|---|---|---|---|
| $T_{IP/IC}$ | local | Trait value distribution is independent of species identity<br><br>there is no internal filtering. | Individual trait values are shuffled within each community | two individuals belonging to a population have more similar trait values than two individuals drawn randomly from the community |
| $T_{I/CIR}$ | Regional. ind | Individual trait value distribution is drawn randomly from the regional pool:<br><br>there is no external filtering acting on individuals. | Individual trait values are shuffled within the regional pool, keeping the number of individuals in each community constant. | Two individuals belonging to a community have more similar trait values than two individuals drawn randomly from the regional pool |
| $T_{PC/PR}$ | Regional. pop | Species mean trait value distribution is drawn randomly from the regional pool<br><br>there is no external filtering acting on species. | Each individual is assigned the mean value of the species; the values are then shuffled within the regional pool, keeping the number of individuals in each community constant. | Two individuals belonging to a community have more similar population based trait values than two individuals drawn randomly from the regional pool while taking abundance into account |

**Table 2.** Fisher's Z-statistic and associated p-values for a comparison of the slopes of CWM1 (population mean trait values weighted by local abundance) and CWM2 (regional species mean trait values weighted by local abundance) for each *Elevation*-Trait regression (Main; Figure 2). The traits used are: body mass (BM), wing loading (WL) and wing aspect ratio (AR).

| Trait | z-statistic | p.value |
|---|---|---|
| **BM** (CWM1 v/s CWM2) | 0.490 | 0.31 |
| **WL** (CWM1 v/s CWM2) | -0.304 | 0.38 |
| **AR** (CWM1 v/s CWM2) | 0.177 | 0.43 |

**Table 3.** Fisher's Z-statistic and associated p-values for a comparison of the slopes of three *Environment* distance-Trait regressions (Main; Figure 2): body mass (BM), wing loading (WL), wing aspect ratio (AR).

| Trait pair | z-statistic | p.value |
|---|---|---|
| **BM overlap – WL overlap** | -1.188 | 0.12 |
| **WL overlap – AR overlap** | 0.167 | 0.43 |
| **AR overlap – BM overlap** | 1.747 | < 0.05* |

**Table 4.** Regression parameters of trait overlap between adjacent elevational communities and their mean elevation. The three traits used are: body mass (BM), wing loading (WL), and wing aspect ratio (AR).

| Model | Intercept | SE_Intercept | Slope | SE_Slope | R.squared | p.value |
|---|---|---|---|---|---|---|
| **BM overlap ~ Elevation** | 0.85 | 0.03 | $2 \times 10^{-6}$ | $2 \times 10^{-5}$ | -0.10 | 0.92 |
| **WL overlap ~ Elevation** | 0.81 | 0.07 | $4 \times 10^{-5}$ | $4 \times 10^{-5}$ | -0.01 | 0.38 |
| **AR overlap ~ Elevation** | 0.88 | 0.02 | $7 \times 10^{-6}$ | $0.1 \times 10^{-6}$ | -0.06 | 0.54 |

**Table 5.** Observed T-statistics for elevational communities of hawkmoths.
The traits used are: Body Mass (BM), Wing Loading (WL) and Aspect Ratio (AR)

| | **TIP/IC** | | | **TIC/IR** | | | **TPC/PR** | | |
|---|---|---|---|---|---|---|---|---|---|
| | **BM** | **WL** | **AR** | **BM** | **WL** | **AR** | **BM** | **WL** | **AR** |
| **E0200** | 0.26 | 0.40 | 0.76 | 0.84 | 1.71 | 0.62 | 0.73 | 1.14 | 0.49 |
| **E0500** | 0.25 | 0.42 | 0.76 | 0.71 | 0.89 | 1.05 | 0.82 | 1.10 | 0.92 |
| **E0700** | 0.23 | 0.65 | 0.53 | 0.95 | 1.07 | 1.47 | 0.77 | 0.87 | 1.34 |
| **E0900** | 0.33 | 0.36 | 0.54 | 0.74 | 1.24 | 0.86 | 0.76 | 0.99 | 0.54 |
| **E1100** | 0.22 | 0.69 | 0.70 | 1.15 | 0.88 | 1.05 | 1.28 | 1.03 | 1.67 |
| **E1300** | 0.18 | 0.61 | 0.47 | 1.18 | 0.54 | 1.11 | 1.13 | 0.40 | 1.76 |
| **E1500** | 0.29 | 0.48 | 0.48 | 0.93 | 0.84 | 0.76 | 1.18 | 1.15 | 0.60 |
| **E1700** | 0.10 | 0.36 | 0.30 | 1.52 | 0.95 | 1.27 | 1.43 | 1.09 | 1.65 |
| **E1900** | 0.10 | 0.58 | 0.38 | 1.27 | 0.69 | 1.02 | 1.08 | 0.98 | 1.09 |
| **E2100** | 0.20 | 0.82 | 0.53 | 1.14 | 0.65 | 1.00 | 1.12 | 0.77 | 0.61 |
| **E2300** | 0.26 | 0.57 | 0.40 | 0.98 | 0.59 | 0.98 | 1.13 | 0.77 | 0.86 |
| **E2500** | 0.25 | 0.59 | 0.55 | 0.70 | 0.69 | 0.78 | 0.88 | 0.71 | 0.59 |
| **E2700** | 0.21 | 0.70 | 0.62 | 0.77 | 0.71 | 1.01 | 0.89 | 1.10 | 1.08 |

**Table 6. Standard Effect Size for the metric $T_{IP/IC}$ for elevational communities of hawkmoths.**
The columns are Elevational community, SES, 2.75% of the null distribution (Low), 97.5% of the null distribution (High). The calculations were made separately for body mass (BM), wing loading (WL) and wing aspect ratio (AR).

| | **SES** | | | **Null Low** | | | **Null High** | | |
|---|---|---|---|---|---|---|---|---|---|
| | **BM** | **WL** | **AR** | **BM** | **WL** | **AR** | **BM** | **WL** | **AR** |
| **E0200** | -5.98 | -4.82 | -1.55 | -1.75 | -1.74 | -1.46 | 2.14 | 2.29 | 2.66 |
| **E0500** | -4.54 | -3.42 | -1.24 | -1.66 | -1.65 | -1.53 | 2.26 | 2.30 | 2.38 |
| **E0700** | -5.17 | -2.35 | -2.74 | -1.54 | -1.51 | -1.53 | 2.24 | 2.25 | 2.45 |
| **E0900** | -4.06 | -4.06 | -2.55 | -1.71 | -1.65 | -1.64 | 2.22 | 2.18 | 2.38 |
| **E1100** | -5.66 | -2.16 | -1.91 | -1.68 | -1.60 | -1.63 | 2.16 | 2.23 | 2.43 |
| **E1300** | -5.56 | -2.54 | -3.00 | -1.60 | -1.61 | -1.49 | 2.41 | 2.34 | 2.25 |
| **E1500** | -4.12 | -2.80 | -2.63 | -1.64 | -1.56 | -1.63 | 2.26 | 2.24 | 2.38 |
| **E1700** | -4.33 | -3.12 | -2.86 | -1.68 | -1.77 | -1.50 | 2.16 | 2.08 | 2.32 |
| **E1900** | -6.30 | -2.73 | -3.37 | -1.63 | -1.55 | -1.53 | 2.16 | 2.24 | 2.14 |
| **E2100** | -4.44 | -1.04 | -2.21 | -1.63 | -1.61 | -1.40 | 2.40 | 2.39 | 2.36 |
| **E2300** | -3.89 | -2.23 | -2.78 | -1.37 | -1.33 | -1.26 | 2.53 | 2.50 | 2.67 |
| **E2500** | -3.37 | -1.97 | -1.79 | -1.36 | -1.37 | -1.29 | 2.38 | 2.54 | 2.45 |
| **E2700** | -3.79 | -1.48 | -1.56 | -1.64 | -1.58 | -1.50 | 2.20 | 2.42 | 2.42 |

**Table 7. Standard Effect Size for the metric $T_{IC/IR}$ for elevational communities of hawkmoths.**

The columns are Elevational community, SES, 2.75% of the null distribution (Low), 97.5% of the null distribution (High). The calculations were made separately for body mass (BM), wing loading (WL) and wing aspect ratio (AR).

| | BM | WL | AR | BM_low | WL_low | AR_low | BM_high | WL_high | AR_high |
|---|---|---|---|---|---|---|---|---|---|
| **E0200** | -2.20 | 8.81 | -3.76 | -1.84 | -1.97 | -1.84 | 2.01 | 2.00 | 2.15 |
| **E0500** | -3.29 | -1.15 | 0.45 | -1.78 | -1.82 | -1.82 | 2.02 | 2.12 | 2.15 |
| **E0700** | -0.65 | 0.64 | 3.76 | -1.85 | -1.82 | -1.74 | 1.87 | 2.08 | 2.18 |
| **E0900** | -2.52 | 2.02 | -1.02 | -1.73 | -1.84 | -1.68 | 2.03 | 2.08 | 2.20 |
| **E1100** | 1.62 | -1.07 | 0.31 | -1.92 | -1.88 | -1.75 | 2.01 | 2.01 | 2.13 |
| **E1300** | 1.57 | -3.93 | 0.78 | -1.84 | -1.86 | -1.75 | 2.01 | 2.06 | 2.26 |
| **E1500** | -0.57 | -1.24 | -1.56 | -1.84 | -1.73 | -1.71 | 2.07 | 2.16 | 2.16 |
| **E1700** | 3.24 | -0.25 | 1.34 | -1.76 | -1.83 | -1.62 | 2.20 | 2.08 | 2.35 |
| **E1900** | 3.17 | -3.15 | 0.15 | -1.85 | -1.98 | -1.74 | 1.88 | 2.11 | 2.23 |
| **E2100** | 1.30 | -2.89 | 0.08 | -1.94 | -1.84 | -1.72 | 2.07 | 2.09 | 2.16 |
| **E2300** | -0.24 | -4.03 | -0.15 | -1.86 | -1.93 | -1.70 | 2.11 | 1.95 | 2.05 |
| **E2500** | -3.15 | -2.89 | -1.67 | -1.95 | -1.87 | -1.73 | 2.00 | 2.07 | 2.17 |
| **E2700** | -2.59 | -2.94 | 0.10 | -1.85 | -1.94 | -1.73 | 2.06 | 2.06 | 2.05 |

**Table 8. Standard Effect Size for the metric $T_{PC/PR}$ for elevational communities of hawkmoths.**

The columns are Elevational community, SES, 2.75% of the null distribution (Low), 97.5% of the null distribution (High). The calculations were made separately for body mass (BM), wing loading (WL) and wing aspect ratio (AR).

| | BM | WL | AR | BM_low | WL_low | AR_low | BM_high | WL_high | AR_high |
|---|---|---|---|---|---|---|---|---|---|
| **E0200** | -0.55 | 0.69 | -1.14 | -1.59 | -1.43 | -1.28 | 2.17 | 2.34 | 3.01 |
| **E0500** | -0.34 | 0.39 | -0.11 | -1.53 | -1.43 | -1.26 | 2.21 | 2.36 | 2.90 |
| **E0700** | -0.01 | 0.21 | 1.33 | -1.41 | -1.32 | -1.17 | 2.22 | 2.48 | 2.62 |
| **E0900** | -0.58 | 0.04 | -1.10 | -1.58 | -1.42 | -1.30 | 2.34 | 2.45 | 2.69 |
| **E1100** | 0.72 | 0.07 | 1.62 | -1.59 | -1.52 | -1.28 | 2.09 | 2.23 | 2.79 |
| **E1300** | 0.29 | -1.20 | 1.62 | -1.53 | -1.31 | -1.18 | 2.30 | 2.43 | 2.54 |
| **E1500** | 0.13 | 0.06 | -1.23 | -1.60 | -1.54 | -1.35 | 2.25 | 2.32 | 2.76 |
| **E1700** | 0.54 | -0.11 | 0.88 | -1.50 | -1.36 | -1.22 | 2.25 | 2.60 | 2.86 |
| **E1900** | 0.16 | -0.12 | 0.23 | -1.62 | -1.53 | -1.28 | 2.23 | 2.31 | 2.81 |
| **E2100** | 0.30 | -0.42 | -0.81 | -1.54 | -1.24 | -1.19 | 2.23 | 2.52 | 2.94 |
| **E2300** | 0.26 | -0.45 | -0.31 | -1.51 | -1.35 | -1.15 | 2.40 | 2.61 | 2.83 |
| **E2500** | -0.36 | -0.56 | -0.87 | -1.39 | -1.24 | -1.23 | 2.33 | 2.57 | 2.95 |
| **E2700** | -0.47 | 0.05 | 0.01 | -1.52 | -1.43 | -1.24 | 2.23 | 2.54 | 2.96 |

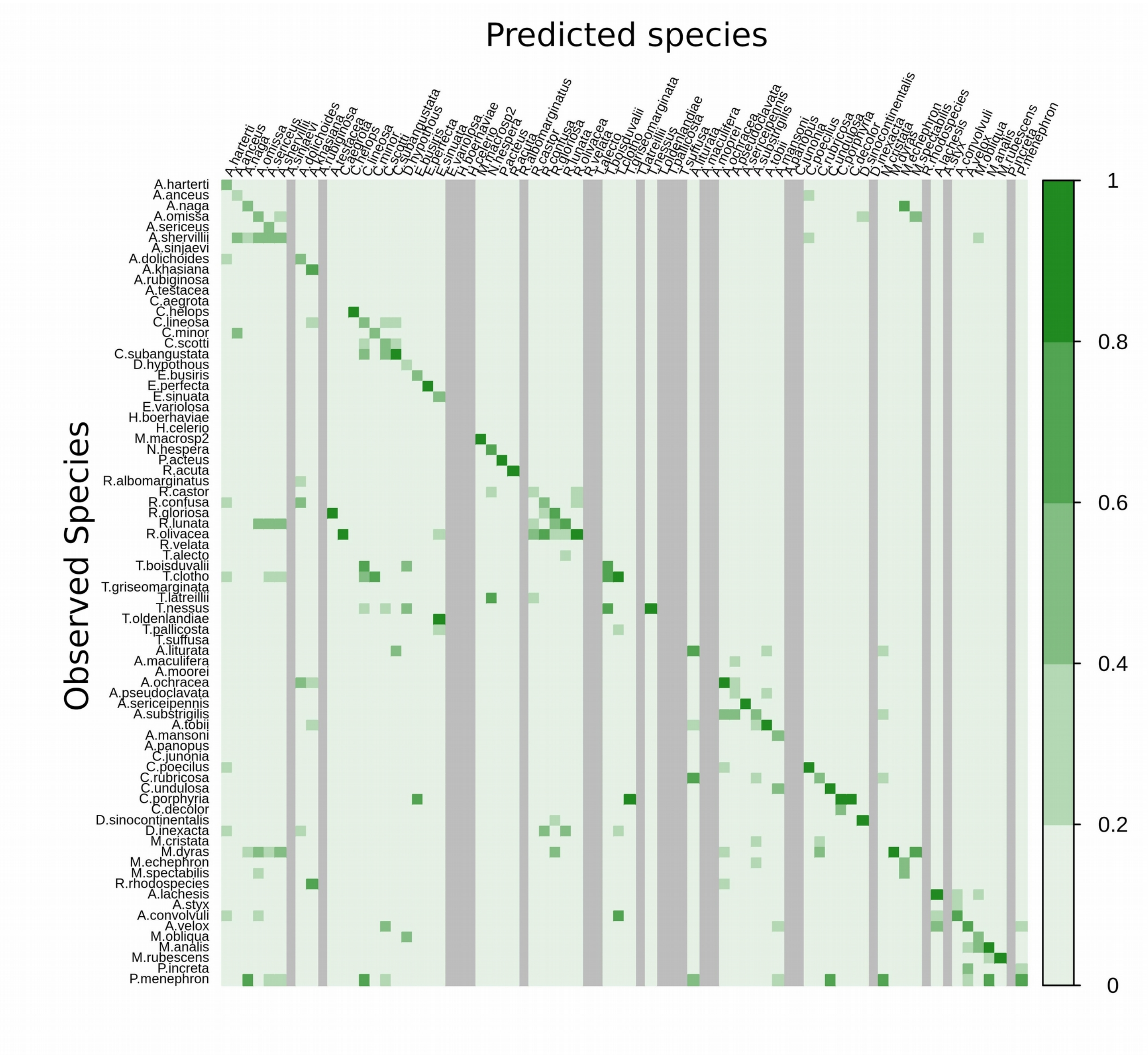


**Figure 1.** Probability of assigning a hawkmoth individual to its correct species from Linear Discriminant Analysis (LDA) using the primary traits of body-length, thorax width, wing length and wing breadth. Values greater than 0 outside the diagonal indicate errors in attributing the individual to it's correct species. Grey color indicates failure of LDA to assign individuals to the given species.

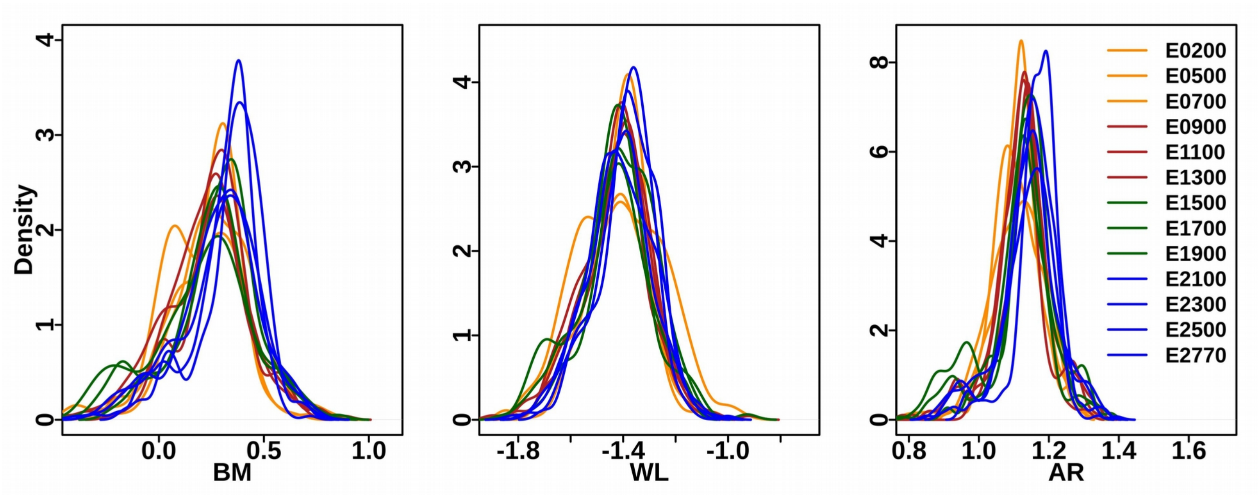


**Figure 2. Kernel density plots of traits of each elevational community.**
Each color coded curve represents the normalized density distribution for an elevational community. Low elevations (200, 500, 700 & 900 m) are shown in red, mid-elevations (1100, 1300, 1500 & 1700 m) in green and high elevations (2100 m and above) in blue. The different traits plotted are body mass (BM), wing loading (WL) and aspect ratio (AR).

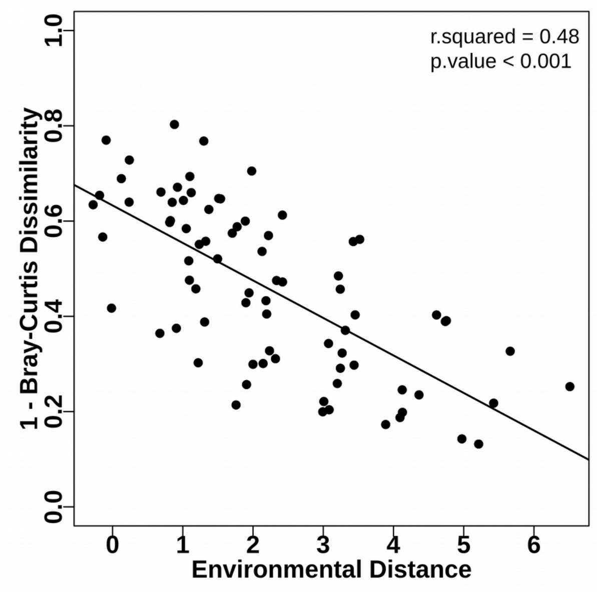


**Figure 3. Relationship of species turnover and *Environment*al distance.**
The species turnover was characterized by the Bray-Curtis dissimilarity index between all pairs of communities.

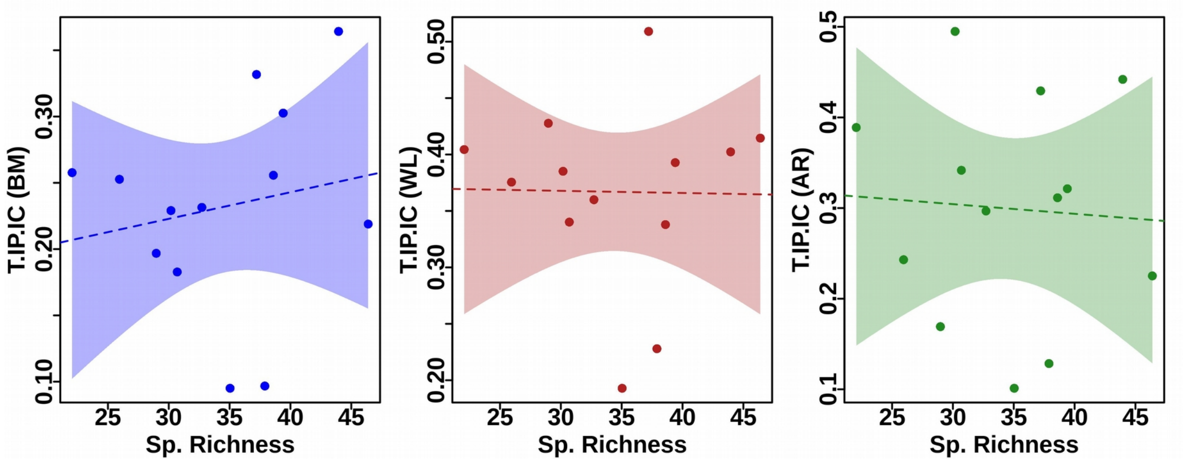


**Figure 4.** Relationship between observed T-statistic $T_{IP/IC}$ and rarefied species richness The traits used are body mass (BM), wing loading (WL) and wing aspect ratio (AR). None of the traits exhibited a significant relationship between the two variables.
